# Distinct myofibre domains of the human myotendinous junction revealed by single nucleus RNA-seq

**DOI:** 10.1101/2022.12.16.519020

**Authors:** Anders Karlsen, Ching-Yan Chloé Yeung, Peter Schjerling, Linda Denz, Christian Hoegsbjerg, Jens R. Jakobsen, Michael R. Krogsgaard, Manuel Koch, Stefano Schiaffino, Michael Kjaer, Abigail L. Mackey

## Abstract

The myotendinous junction (MTJ) is a specialized domain of the multinucleated myofibre, faced with the challenge of maintaining robust cell-matrix contact with the tendon under high mechanical stress and strain. Here, we profiled 24,161 nuclei in semitendinosus muscle-tendon samples from 3 healthy males by single nucleus RNA-sequencing (snRNA-seq), alongside spatial transcriptomics, to gain insight into the genes characterizing this specialization in humans. We identified a cluster of MTJ myonuclei, represented by 47 enriched transcripts, of which the presence of *ABI3BP, ABLIM1, ADAMTSL1, BICD1, CPM, FHOD3, FRAS1* and *FREM2* was confirmed at the MTJ at the protein level by immunofluorescence. Four distinct subclusters of MTJ myonuclei were apparent and segregated into two COL22A1-expressing subclusters and two lacking COL22A1 but with a clear fibre type profile expressing *MYH7* or *MYH1/2*. Our findings reveal distinct myonuclei profiles of the human MTJ, a weak link in the musculoskeletal system, which is selectively affected in pathological conditions, from muscle strains to muscular dystrophies.

## Introduction

The myotendinous junction (MTJ) is highly specialized in both structure and molecular composition, designed to meet the challenge of linking two distinctive tissues and withstanding substantial forces (Tidball 1991; Kojima et al. 2008; Knudsen et al. 2015). As the interface between muscle and the matrix-rich tendon, the MTJ is characterized by enrichment of certain proteins, the most prominent being collagen XXII (Koch et al. 2004). Collagen XXII is found in the basement membrane only at the myofibre tip and is seen in tendon attachment sites of muscles with diverse functions, such as limb and intercostal skeletal muscles, and the papillary muscles of the heart where they attach to the chordae tendineae (Koch et al. 2004). Recently by proteomics, we identified 112 proteins enriched at the human semitendinosus MTJ when compared to neighbouring muscle and tendon tissue (Karlsen et al. 2022), but due to the nature of the MTJ as a muscle-tendon composite, this approach did not allow for resolving the origin of these proteins. Single-nucleus RNA-sequencing (snRNA-seq) has led to major progress in the understanding of the transcriptional diversity in the multinucleated skeletal muscle fibre, and recent studies in mice consistently found a cluster of transcriptionally distinct myonuclei, presumably encoding genes for MTJ proteins (Dos Santos et al. 2020; Chemello et al. 2020; Kim et al. 2020; Petrany et al. 2020; Wen et al. 2021).

The only other known transcriptionally distinct domain of the myofibre is the neuromuscular junction (NMJ). The main portion of the myofibre contains thousands of myonuclei, each believed to manage their own domain, but under a common transcriptional programme, to provide mRNA for the major structural and metabolic functions of the myofibre. While the NMJ has received extensive attention, much less focus has been directed to the MTJ, although this myofibre domain is selectively affected in different pathological clinical conditions, from muscle strain injuries (Grange et al. 2022) to muscular dystrophies (Chemello et al. 2020). Interestingly, in mice with skeletal muscle-specific disruption of dystroglycan, dystroglycan staining was curiously preserved in the sarcolemma at the NMJ and MTJ (Cohn et al. 2002), further supporting the notion of segregated domains in the myofibre. Furthermore, muscle ends also appear to be the site of disease initiation (Heskamp et al. 2022), the reason for which remains unknown. Therefore, elucidation of the basic biological regulation of the MTJ domain is important in understanding the progression and treatment of a wide range of unsolved pathologies.

Here, we performed the first snRNA-seq analysis of the human muscle-tendon unit, revealing MTJ-enriched molecules and sub-specialization of the myonuclei at the tip of the myofibre. Published snRNA-seq analysis of human skeletal muscle is currently limited to tissue collected post-mortem, often with substantial ischemic time and limited characterisation of donors (Eraslan et al. 2022). To circumvent this issue, we collected surgical waste muscle-tendon tissue from three well-characterized healthy individuals, with the use of strict exclusion and inclusion criteria, and tissue was frozen within 40 minutes of surgical extraction. Freezing the tissue also facilitated running the samples in the same batch. Other strengths of our approach include the avoidance of enzymatic tissue digestion to preserve the transcriptional state more than with single cell RNA-seq. Lastly, we applied a conservative statistical approach requiring DEGs (Differentially Expressed Genes) to be significantly enriched in a cluster in each of the 3 individuals, thus accounting for biological variation. Therefore, in addition to our focus on the MTJ-myonuclei, our data also provide high quality profiling of nuclei in healthy human tendon and muscle tissue. Spatial transcriptomics and immunofluorescence supported our snRNA-seq data and together advance the understanding of the MTJ myonuclei and the mononuclear cells present in healthy human muscle and tendon tissue.

## Results

### snRNA-seq profile of the human myotendinous junction in the context of muscle-tendon tissue

From three human samples (Figure 1), cDNA libraries of a total of 25,556 nuclei were prepared with the 10x Chromium system, sequenced using droplet based single nuclei RNA-seq and subjected to bioinformatical analysis (Table 1). Of these nuclei, 3-7% were identified as doublets and removed, resulting in 6,575, 10,140, and 7,409 nuclei from the three participants for the downstream analyses (Table 1). The three data sets were integrated and subjected to unbiased clustering, resulting in 17 clusters (Figure 2A). The clusters included all the expected cell types in tendon and skeletal muscle (Figure 2A), and all clusters contained nuclei from all three samples (Supplementary Figure S1A). A list of DEGs for each cluster can be found in Supplementary Data S1, and a complete overview of gene-symbol abbreviations, Ensemble gene IDs and gene names is provided in Supplementary Table S1.

**Table 1:** Overview of number of nuclei at all stages of the snRNA-seq process

|  | Sample S1 | Sample S3 | Sample S4 |
| --- | --- | --- | --- |
| Nuclei loaded | $1.7 \cdot 10^4$ | $1.6 \cdot 10^4$ | $1.6 \cdot 10^4$ |
| Nuclei sequenced | 6740 | 10965 | 7851 |
| Mean UMI counts | 1817 | 4557 | 6012 |
| Mean genes per nuclei | 817 | 1715 | 2020 |
| Identified doublets | 165 (2.4%) | 825 (7.5%) | 442 (5.6%) |
| Final number of nuclei | 6575 | 10140 | 7409 |

*Abbreviations: UMI - Unique Molecular Identifier*

**Figure 1.**
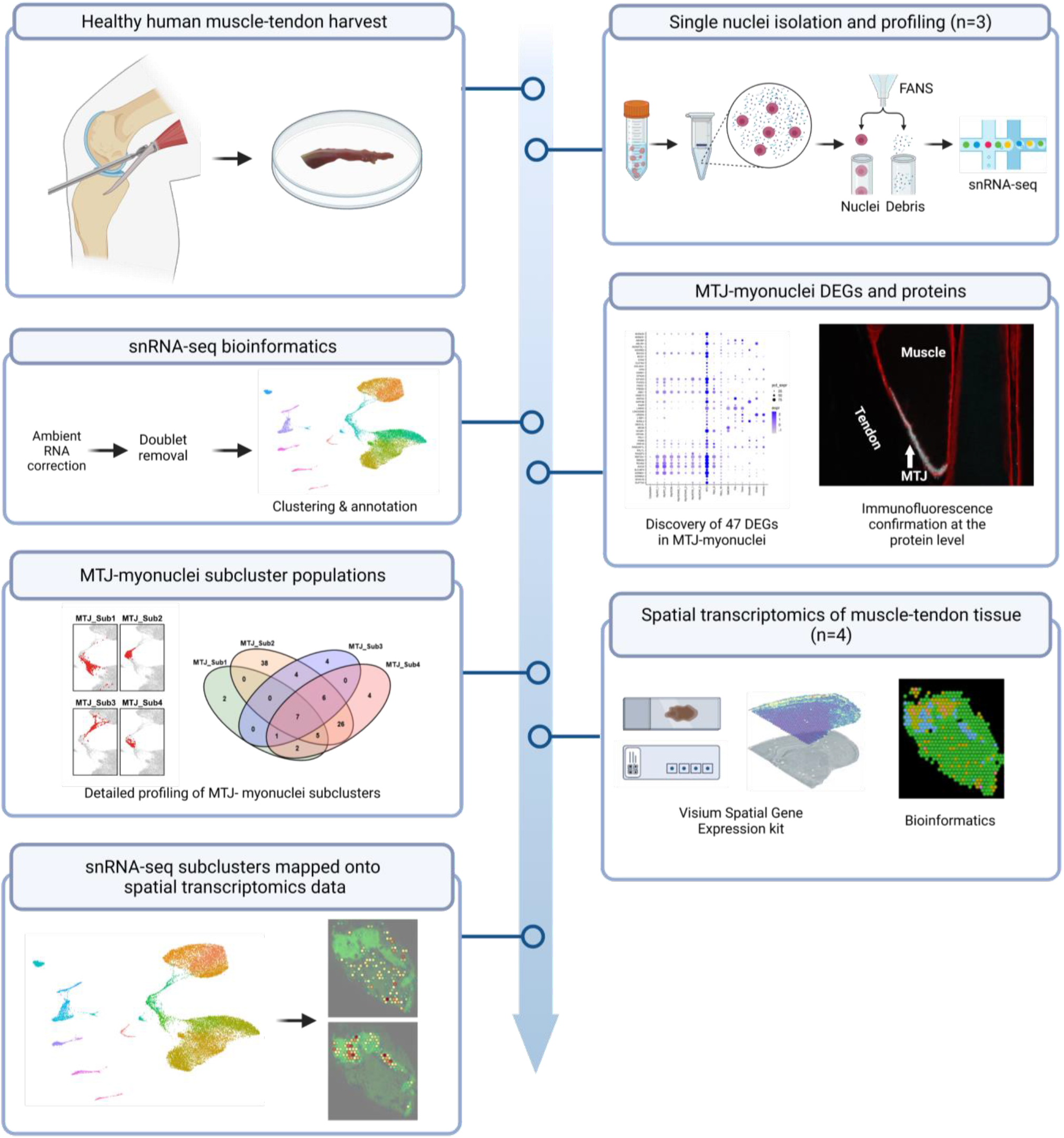
Workflow. Semi-tendinosis muscle-tendon tissue collected from patients during ACL surgery, placed on ice within 8 minutes, cut in small pieces and snap frozen before stored at −80°C. Frozen samples were then cut in smaller pieces in a cryochamber (−20°C) and stored at −80°C, followed by nuclei extraction and Fluorescence-Activated Nuclei Sorting (FANS). Libraries were constructed with the 10x Chromium platform followed by sequencing. Bioinformatics resulted in 17 nuclei clusters with 47 differentially expressed genes (DEGs) in the MTJ-myonuclei cluster, followed by immunofluorescence (IF) confirmation at the protein level. Analysis of 4 MTJ-subclusters and mapping of snRNA-seq nuclei clusters onto spatial transcriptomics data obtained with the Visium Spatial Gene Expression kit (10x Genomics). Created with BioRender.com.

**Figure 2.**
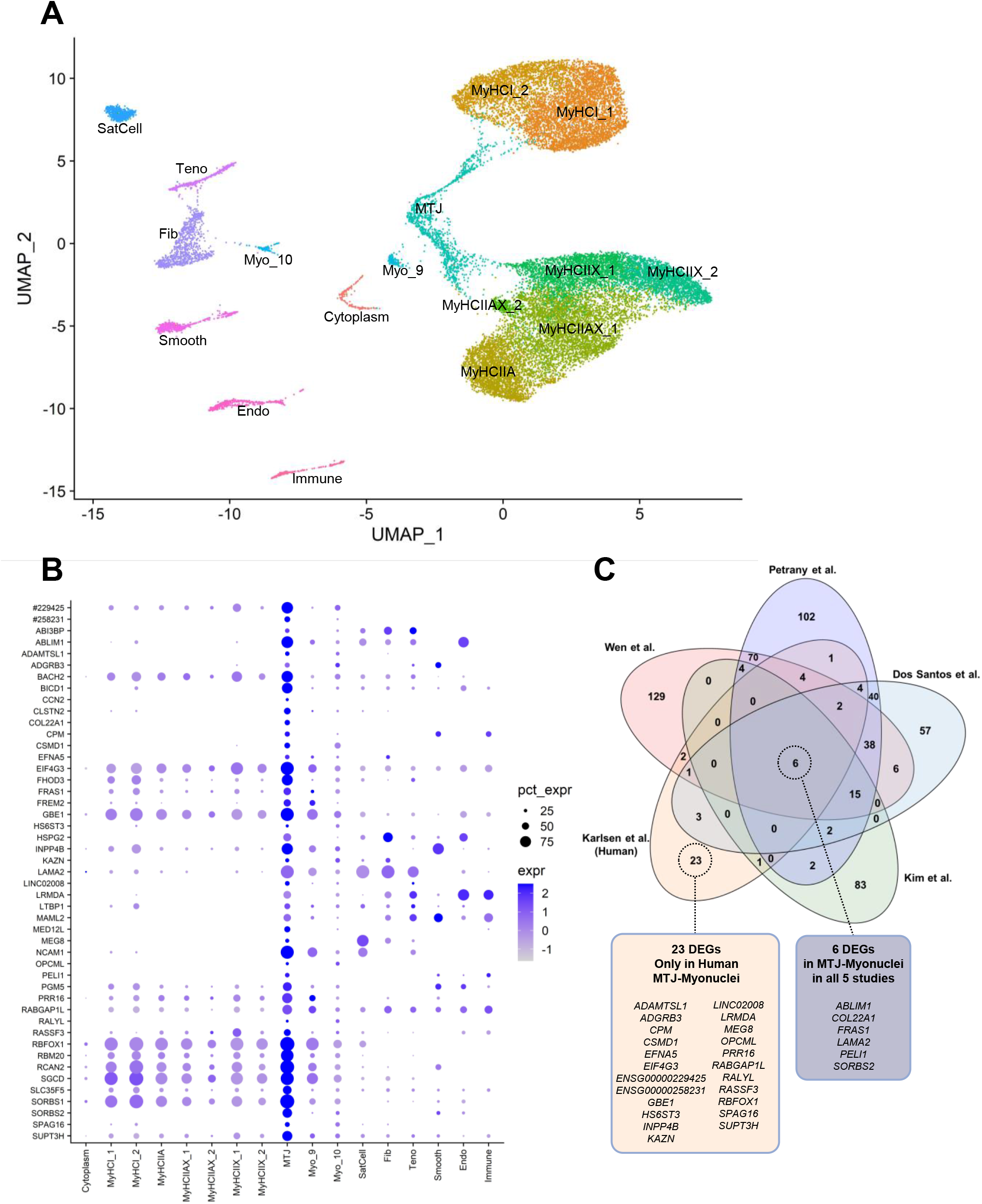
snRNA-seq derived DEGs of the human myotendinous junction. **A:** snRNA-seq UMAP illustrating the 17 nuclei clusters in the snRNA-seq analysis of human muscle-tendon tissue (n=3). **B:** Dot-plot of the expression profile of the 47 differentially enriched genes (DEGs) in the MTJ-myonuclei cluster. **C:** Venn diagram comparing MTJ-myonuclei enriched transcripts in mice (Dos Santos et al. 2020; Kim et al. 2020; Petrany et al. 2020; Wen et al. 2021) and in human (Karlsen et al.; this study).

Six non-myonuclei clusters were assigned through known marker genes for satellite cells (*PAX7*), fibroblasts (*COL1A2, COL3A1, FAP, DCN, TCF7L2* and PDGFRA), tenocytes (*COMP* and *MKX*), smooth muscle cells (*ACTA2, MYH11* and *RGS5*), endothelial cells (*VWF* and *PECAM1*) and immune cells (*CD163* and *F13A1*) (Figure 2A, Supplementary Figure S1B, Supplementary Data S1). The remaining 11 clusters were myonuclei, based on titin (TTN) expression, and clustered according to myofibre type. We detected two type I myosin clusters (*MYH7*), one type IIA myosin cluster (*MYH2*), two type IIX myosin clusters (*MYH1*), two type IIAX myosin clusters (*MYH1, MYH2*) and one MTJ-myonuclei cluster (*COL22A1, NCAM1, SORBS2*) expressing all three major myosin heavy chains (*MYH1, MYH2* and *MYH7*) (Figure 2A, Supplementary Figure S1B). Two myofibre clusters (Myo_9, Myo_10) were not assigned an identity due to the lack of clear markers. The Myo_9 cluster was mainly found in one sample (Supplementary Figure S1A) and is a NMJ cluster candidate since it was the only cluster containing genes previously identified in NMJ myonuclei clusters, for example, the acetylcholine receptor *CHRNA1* (Kim et al. 2020; Petrany et al. 2020), along with *RAPH1* (Dos Santos et al. 2020; Chemello et al. 2020; Petrany et al. 2020), *ITGA9* (Dos Santos et al. 2020; Chemello et al. 2020; Petrany et al. 2020), and *PDGFC* (Dos Santos et al. 2020; Petrany et al. 2020) (Supplementary Data S1). The second unassigned cluster, Myo_10, was not fibre type specific (Supplementary Figure S1B). The last myonuclei cluster, the “Cytoplasm” cluster, contained high levels of mitochondrial RNAs, e.g. *MT-CO1* (Supplementary Figure S1B). A large proportion of these “nuclei” did not express the nuclei marker *MALAT1*, indicating that this cluster could represent cytoplasmic RNA, and not nuclei (Supplementary Figure S1B). However, a similar Cytoplasm cluster has been observed by others and it is currently under debate whether this cluster is an artefact or not (Tucker et al. 2020; Eraslan et al. 2022).

The main purpose of this study was to examine the MTJ-myonuclei in human, which in our data set was characterized by 47 DEGs. The top 6 most differentially expressed marker genes were *SORBS2, BICD1, COL22A1, NCAM1, MED12L* and *ABLIM1* (Figure 2B, Supplementary Data S1). First, we compared these 47 DEGs with previous MTJ-myonuclei data in mice (Figure 2C). We used data from three mouse studies on uninjured muscles (Dos Santos et al. 2020; Petrany et al. 2020; Wen et al. 2021) and one mouse study on a mix of uninjured and regenerating muscles (Kim et al. 2020). Approximately half (23 out of 47) of the DEGs in human MTJ-myonuclei were observed in mouse MTJ-myonuclei (Figure 2C), but only 6 DEGs were found in all 5 datasets (*ABLIM1, COL22A1, FRAS1, LAMA2, PELI1, SORBS2*, Figure 2C, Supplementary Data S2).

### Immunofluorescence confirms snRNAseq DEGs at the MTJ

Next, we used immunofluorescence (IF) to investigate if the DEGs in our human MTJ-myonuclei translated into protein enrichment in human MTJ tissue, and the combined snRNA-seq and IF led to the discovery of 8 MTJ-enriched proteins (Figure 3, Supplementary Figures S2-S3). Most of these MTJ-enriched proteins were localized near the muscle fibre membrane in close proximity to the collagen XXII defined MTJ (FRAS1, FREM2, BICD1, FHOD3, CPM, ADAMTSL1, Figure 3A-F). The IF signal for ABLIM1 was confined to the muscle fibre cytoplasm, and was most intense at the tip of the fibres, gradually fading in the direction of the main portion of the muscle fibre (Figure 3G). The ABI3BP signal was in the tendon and the extracellular matrix between muscle and tendon with a stronger IF signal near the MTJ (Figure 3H). Although ABI3BP was enriched in MTJ-myonuclei, it is possible that the site of *ABI3BP* protein deposition originates from the Fib- and Teno-clusters, where *ABI3BP* transcripts were also detected (Figure 2B, Supplementary Data S1). Notably, these 8 positive IF-targets showed a varying degree of overlap with the DEGs reported in MTJ-myonuclei in mice, as some were found in all 4 mouse datasets (*ABLIM1, FRAS1*); some were only found in one or two of the mouse datasets (ABI3BP, *BICD1, FHOD3, FREM2*), and some were only found in human MTJ-myonuclei (*ADAMTSL1, CPM*) (Figure 2C, Supplementary Data S2), underlining the good agreement as well as diversity between these 5 datasets. In addition to the discovery of new components of the MTJ, the outcomes of our snRNA-seq analysis also demonstrate the robustness of our dataset from 3 subjects and >21,000 profiled nuclei.

**Figure 3.**
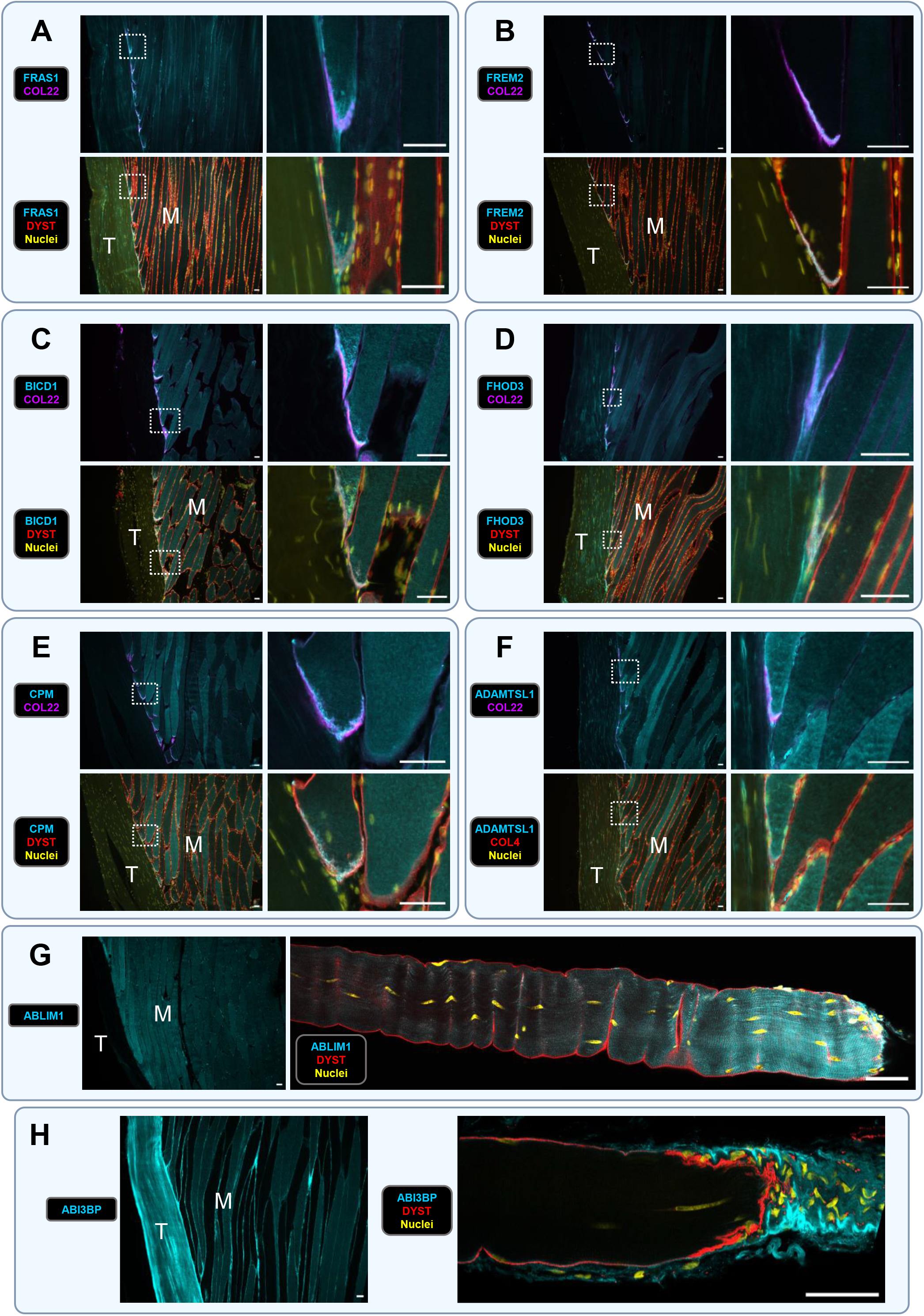
FRAS1, FREM2, BICD1, FHOD3, CPM, ADAMTSL1, ABLIM1 and ABI3BP at the human MTJ. Skeletal muscle fibre region is labelled with “M” and the tendon with “T”. DAPI or Hoechst stained the nuclei and Collagen XXII (COL22) is used as a marker of the myotendinous junction. All scale bars are 50μm. **A-F:** Triple immunofluorescence labelling and widefield microscopy of 10μm thick sections of longitudinally cut human semitendinosus muscle-tendon tissue samples. FRAS1 **(A)**, FREM2 **(B)**, BICD1 **(C)**, FHOD3 **(D)**, CPM **(E)** and ADAMTSL1 **(F)** show strong enrichment at the tip of the muscle fibres in close association with the dystrophin (DYST) stained sarcolemma, or collagen IV (COL4) stained basement membrane, and the MTJ-marker collagen XXII (COL22). The second panel is a higher magnification of the region in the dotted box in the first panel. **G-H:** Panels to the left show widefield microscopy images of ABLIM1 **(G)** and ABI3BP **(H)** in longitudinally cut muscle-tendon tissue sections. ABLIM1 **(G)** stained the cytoplasm strongest near the MTJ, and gradually faded along the length of the muscle fibre. ABI3BP **(H)** staining was strongest in the MTJ-region on the extracellular matrix side of the muscle fibres and in the tendon. The panels to the right show confocal microscopy images of isolated gracilis single fibres. The graded increase in ABLIM1 intensity towards the MTJ in the cytoplasm of the muscle fibre is clearly depicted in **G**, and the extracellular localization of ABI3BP at the tip of the muscle fibre is clearly visible in **H**. Dystrophin (DYST) stained the sarcolemma and Hoechst the nuclei.

### snRNA-seq subcluster analysis reveals distinct myofibre domains

To further explore the transcriptional heterogeneity within the MTJ-myonuclei, we subjected the main clusters to subclustering by repeating the clustering selectively on either MTJ, (MyHCI_1, MyHCI_2), (MyHCIIA, MyHCIIAX_1, MyHCIIAX_2, MyHCIIX_1, MyHCIIX_2), Immune, (Fib, Teno), Smooth, Endo. This gave rise to a total of 30 subclusters including 4 MTJ-myonuclei subclusters; 26 were new subclusters (indicated by ‘Sub’ in the cluster names) and 4 original (Cytoplasm, Myo_9, Myo_10, SatCell) (Figure 4A, Supplementary Figure S4, Supplementary Data S3). To compare cell-cluster markers we include in Supplementary Figure S5 a comparison between our 30 nuclei-subclusters and cell-types or nuclei-types reported in the literature (Harvey et al. 2019; Dos Santos et al. 2020; Kendal et al. 2020; Rubenstein et al. 2020; De Micheli et al. 2020a, b; Barruet et al. 2020; Chemello et al. 2020; He et al. 2020; Kim et al. 2020; Petrany et al. 2020; Perez et al. 2021; Wen et al. 2021; Eraslan et al. 2022; Scripture-Adams et al. 2022; Yan et al. 2022). This showed a good alignment between our clusters compared to snRNA-seq (Perez et al. 2021; Eraslan et al. 2022) and scRNA-seq (Rubenstein et al. 2020; He et al. 2020) in human skeletal muscle, and in snRNA-seq (Dos Santos et al. 2020; Chemello et al. 2020; Wen et al. 2021) and scRNA-seq (De Micheli et al. 2020b) in mice.

**Figure 4.**
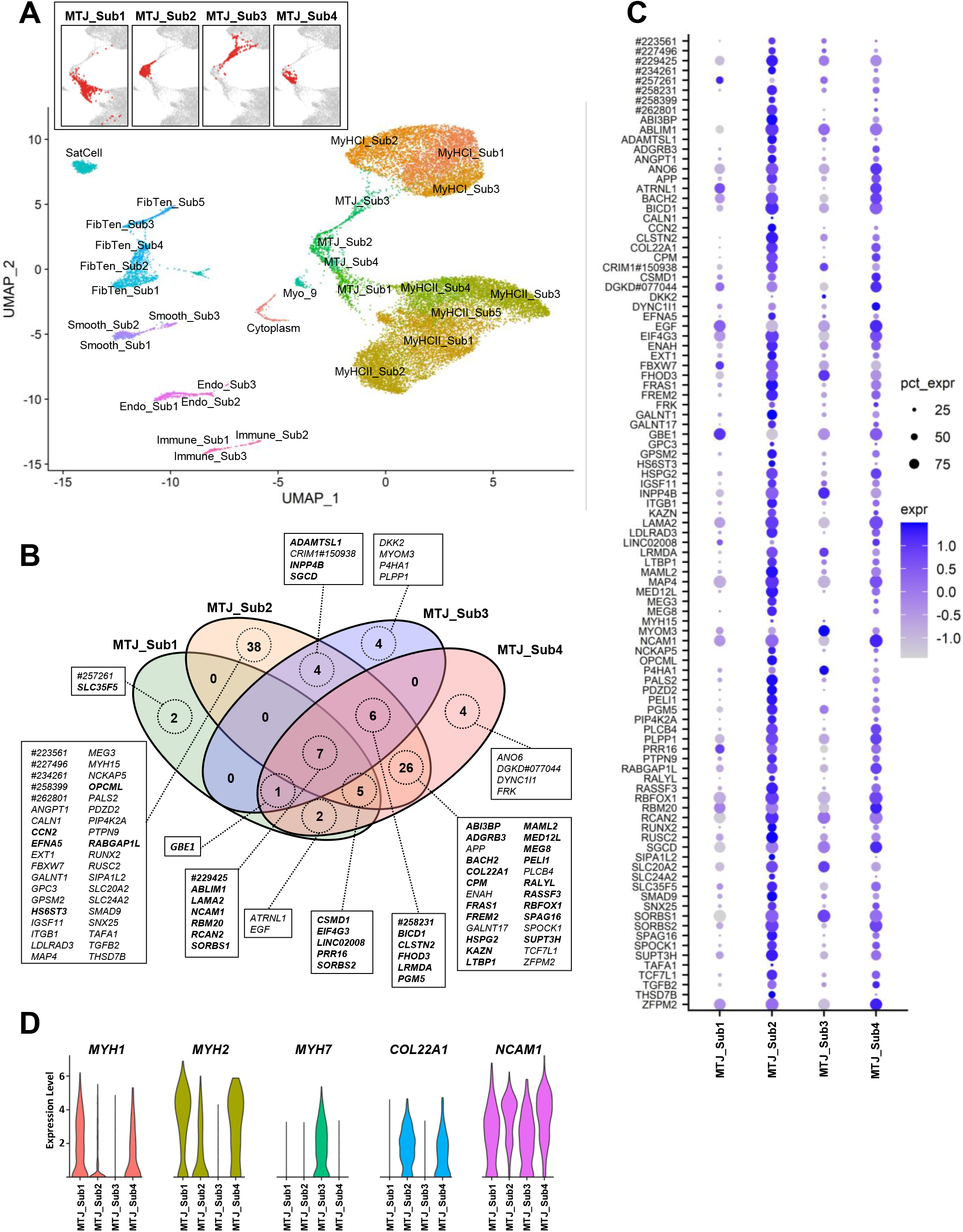
snRNA-seq subcluster profile of the human myotendinous junction. **A:** UMAP illustrating the 30 snRNA-seq nuclei subclusters in human semitendinosus muscle-tendon tissue. In the insert panel the position of nuclei in each of the 4 MTJ-myonuclei subclusters is displayed separately in red colour. **B:** Venn-Diagram showing the overlap of the 99 MTJ DEGs within the 4 MTJ-subclusters. The 47 MTJ-myonuclei DEGs (differentially expressed genes) from the MTJ-myonuclei cluster are highlighted in **bold**. **C:** Dot-plot showing the expression profile of the 99 DEGs in the 4 MTJ-myonuclei subclusters **D:** Violin plot of gene expression levels in the 4 MTJ-myonuclei subclusters. Genes are *MYH1* (myosin heavy chain IIX), MYH2 (myosin heavy chain IIA), *MYH7* (myosin heavy chain I), *COL22A1* (collagen XXII), NCAM1 (neural cell adhesion molecule 1).

The 4 MTJ-myonuclei subclusters (Figure 4A) together expressed 99 DEGs (Figure 4B-C, Supplementary Figure S6, Supplementary Data S3). Of these, 15 coded for proteins belonging to the Matrisome (Naba et al. 2012), divided into: 1) core matrisome genes: collagens (*COL22A1*), extracellular matrix (ECM) glycoproteins (*ABI3BP, FRAS1, LAMA2, LTBP1, CCN2*) and proteoglycans (*HSPG2, SPOCK1*); and 2) matrisome-associated genes: ECM regulators (*ADAMTSL1, P4HA1*), ECM-affiliated proteins (*FREM2, GPC3*), and secreted factors (*ANGPT1, EGF, TGFB2*). The 99 DEGs furthermore encoded for proteins associated with actin assembly (*FHOD3, ENAH, SORBS2*), the microtubule/dynein complex (*BICD1, DYNC1I1, GPSM2, KAZN, NCKAP5*), TGF-β signaling (*SMAD9, SNX25, TGFB2, TCF7L1*) and heparan sulfate chains (*SPOCK1, EXT1, GPC3, HS6ST3*).

Transcriptional heterogeneity between the 4 MTJ-myonuclei subclusters was evident in the form of two *COL22A1*-positive subclusters (*COL22A1*^+^) with predominantly fast-type myosin (MYH1 and/or MYH2, subsequently referred to as *MYH1/2* collectively) expression (MTJ_Sub2 and MTJ_Sub4), and two *COL22A1*-negative subclusters (*COL22A1*^−^) with fibre type specificity (MTJ_Sub1 and MTJ_Sub3), expressing predominantly fast-type (*MYH1/2, MTJ_Sub1*) or slow-type (*MYH7, MTJ_Sub3*) myosin (Figure 4D). The UMAP indicated greatest similarity between MTJ_Sub2 and MTJ_Sub4, overlapping in the central part of the MTJ-cluster (Figure 4A). In addition, these two subclusters had a high prevalence of the DEGs (93 of 99 DEGs in the MTJ_Sub2 and MTJ_Sub4 cluster, Figure 4B), together suggesting that MTJ_Sub2 and MTJ_Sub4 constitute the population of myonuclei residing at the MTJ-domain, defined by the enrichment of collagen XXII protein. Notably, neither of these COL22A1 subclusters had a high expression of *MYH7* (Figure 4D). In contrast, MTJ_Sub1 and MTJ_Sub3 seemed to be less MTJ-specific (*COL22A1*^−^), and together expressed only 31 of the 99 DEGs (Figure 4B, Supplementary Data S3). In the UMAP these subclusters extended from the central part of the MTJ-cluster towards the MyHCII cluster or the MyHCI cluster (Figure 4A), in agreement with their differential expression of fast (*MYH1/2*) and slow myosin (*MYH7*), respectively (Figure 4D).

### Transitional COL22A1 myofibre domain

Based on these findings we hypothesized that MTJ_Sub1 and MTJ_Sub3 belong to a population of myonuclei residing in a transitional MTJ-domain, located between the collagen XXII-rich MTJ-domain and the main portion of the muscle fibre. Accordingly, we define the transitional MTJ-domain as a region of the muscle fibre without enrichment of the collagen XXII protein, but with high expression of other proteins also enriched in the MTJ-domain and not in the main portion of the muscle fibre. The only protein with this IF-staining pattern was ABLIM1 (Figure 3G, Supplementary Figure S3D), demonstrating a similar staining pattern to NCAM1 at the MTJ (Rieger et al. 1985; Daniloff et al. 1989; Jakobsen et al. 2018, 2021; Karlsen et al. 2022). In line with this staining pattern, *ABLIM1* and *NCAM1* were among the few DEGs enriched in all 4 MTJ-subclusters (Figure 4B, Supplementary Data S3). To further explore these observations, we determined the proportion of nuclei expressing *ABLIM1* and/or *NCAM1* in all 20,314 myonuclei in the MyHC and MTJ-subclusters (Supplementary Data S4). This analysis showed a marked increase in the proportion of nuclei expressing *ABLIM1* and/or *NCAM1* from the myonuclei in the main portion of the fibre (7.3%) to the transitional MTJ_Sub1/MTJ_Sub3 (83.6%), with the peak seen in MTJ-specific MTJ_Sub2/MTJ_Sub4 (97.1%). These observations are schematically illustrated in Figure 5A, together with IF images of ABLIM1 and NCAM1 and their gradually increasing staining intensity towards the MTJ (Figure 5C).

**Figure 5.**
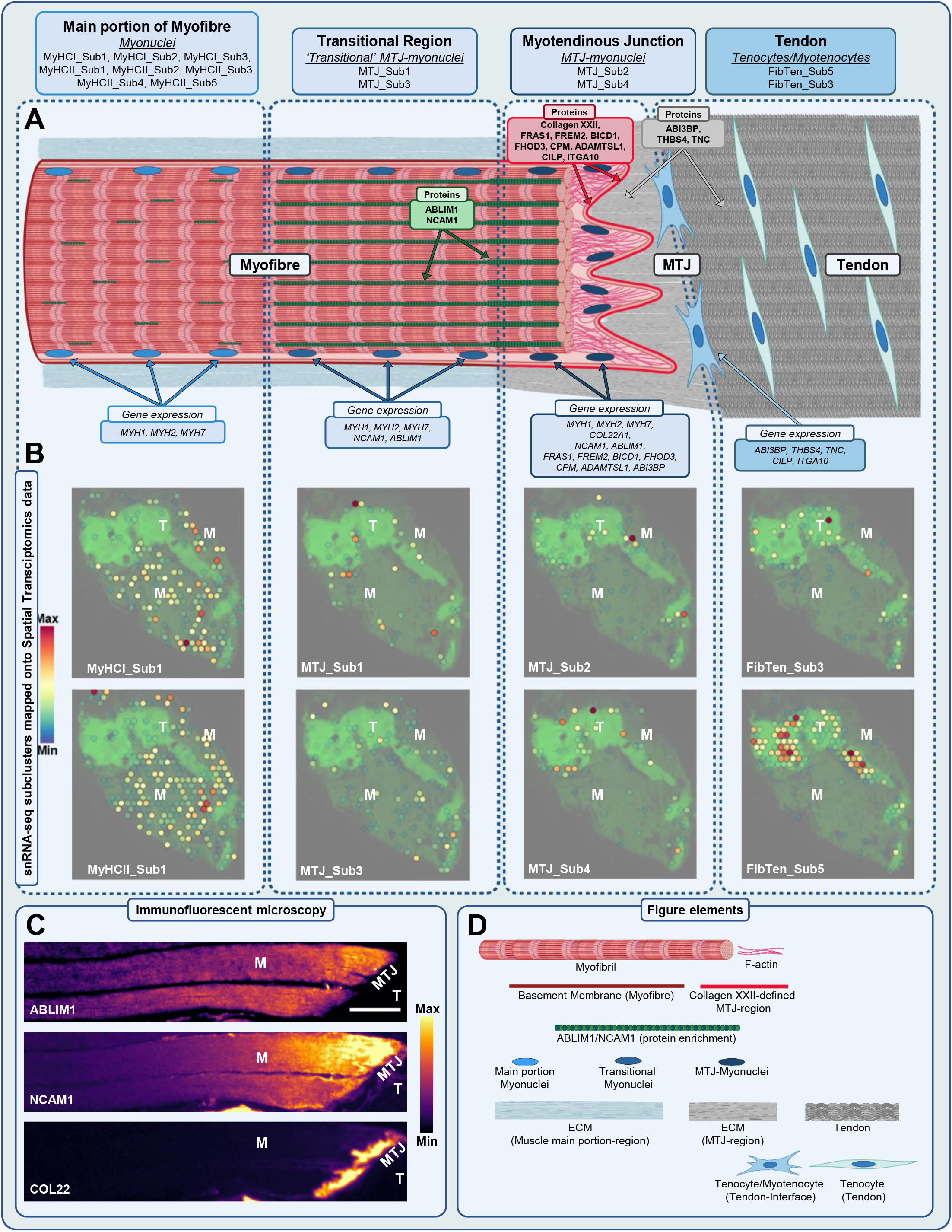
Schematic illustration. Schematic illustration of the main findings. **(A)** The myofibre is divided into three regions; the main portion of the fibre, the transitional region, and the MTJ-region which extends from the muscle fibre to the edge of the tendon. The text boxes link the protein expression from the immunofluorescence, and gene expression from the snRNA-seq data. Spatial transcriptomics mapping of the corresponding snRNA-seq subclusters is presented below **(B). (C)** Single-channel immunofluorescent images from a triple staining for ABLIM1, NCAM1 and COL22 protein, with the signal intensity visualized by a heatmap overlay of the pixel intensity (mpl-inferno, ImageJ). Scale bar is 50 μm. All figure elements are further explained in **(D)**. Note that the MTJ is defined by high expression of collagen XXII protein and expresses several other membrane-associated proteins including FRAS1, FREM2, BICD1, FHOD3, CPM and ADAMTSL1. In the myofibre cytoplasm there is a high abundance of ABLIM1 and NCAM1 protein in the MTJ-region, as well as in the transitional region where the expression of collagen XXII protein is low. This protein expression corresponds to the diverse gene expression profile of the MTJ_Sub2/MTJ_Sub4 (MTJ-myonuclei) vs the MTJ_Sub1/MTJ_Sub3 (transitional myonuclei). In contrast, the abundance of all these MTJ-expressed proteins, including ABLIM1 and NCAM1, is very low in the main portion of the myofibre, corresponding to the gene expression profile of the myonuclei in the main portion of the fibre. The gene expression profile of FibTen_Sub3 included *ABI3BP, THBS4, TNC, CILP* and *ITGA10*, and these associated proteins are highly abundant in the extracellular matrix in the MTJ-region and the tendon (ABI3BP, THBS4, TNC), or in the myofibre membrane at the MTJ (CILP, ITGA10; see Karlsen et. al., 2022). This indicates that FibTen_Sub3 represents MTJ-related “myotenocytes”, in contrast to the tenocyte profile of FibTen_Sub5. Elements of this figure were created with BioRender.com.

### Unexpectedly low abundance of COL22A1^+^ /MYH7^+^ MTJ-myonuclei

The low level of MYH7 expression in COL22A1^+^ MTJ-myonuclei was unexpected because it is generally believed, although not shown, that the collagen XXII protein is enriched at the MTJ in both slow and fast muscle fibres. Therefore, we first confirmed, by IF, the clear presence of collagen XXII protein in slow and fast muscle fibres at the human MTJ (Supplementary Figure S7). Next, we ruled out that *COL22A1^+^ /MYH7*^+^ nuclei were hidden within the non-MTJ-myonuclei clusters, as there were only 30 *COL22A1*^*+*^ nuclei in the non-MTJ-myonuclei subclusters, but none of these were MYH7 positive (Supplementary Data S4). Notably, 19 of these nuclei belonged to the FibTen_Sub3 cluster and were *MYH1/2/7*-negative. We then proceeded with a detailed examination of the *MYH1/MYH2/MYH7* expression in all *COL22A1*^+^ and *COL22A1*^−^ MTJ-myonuclei (Supplementary Data S4), schematically illustrated in Figure 6. Within the total number of 1206 MTJ-myonuclei, 282 were *COL22A1*^*+*^ and 924 were *COL22A1*^−^. The largest proportion (82%) of the *COL22A1*^+^ MTJ-myonuclei was found in MTJ_Sub2/MTJ_Sub4, while only 23.9% of the 924 *COL22A1^−^* MTJ-myonuclei belonged to these MTJ-subclusters. A small fraction of the 282 COL22A1 MTJ-myonuclei expressed MYH7 (16 nuclei = 5.7%), or a mix of *MYH7* and either *MYH1* or *MYH2* (7 nuclei = 2.5%). In contrast, more than half of the *COL22A1* ^+^MTJ-myonuclei expressed at least one of the fast myosins (*MYH1/MYH2*, 150 myonuclei = 53.2%) without expressing MYH7. This resulted in a 9.4 times higher abundance of *COL22A1^+^ /MYH1/2*^+^ vs *COL22A1^+^ /MYH7*^+^ myonuclei (Figure 6, Supplementary Data S4). Due to a larger proportion of MyHCII vs MyHCI myonuclei, we computed correction factors for this and other potential confounders (see the methods section for correction factors), resulting in a 4.2 times higher abundance of *COL22A1*^+^ */MYH1/2*^+^ vs *COL22A1*^+^ */MYH7*^+^ myonuclei after correction, suggesting that the underlying explanation is biological.

**Figure 6.**
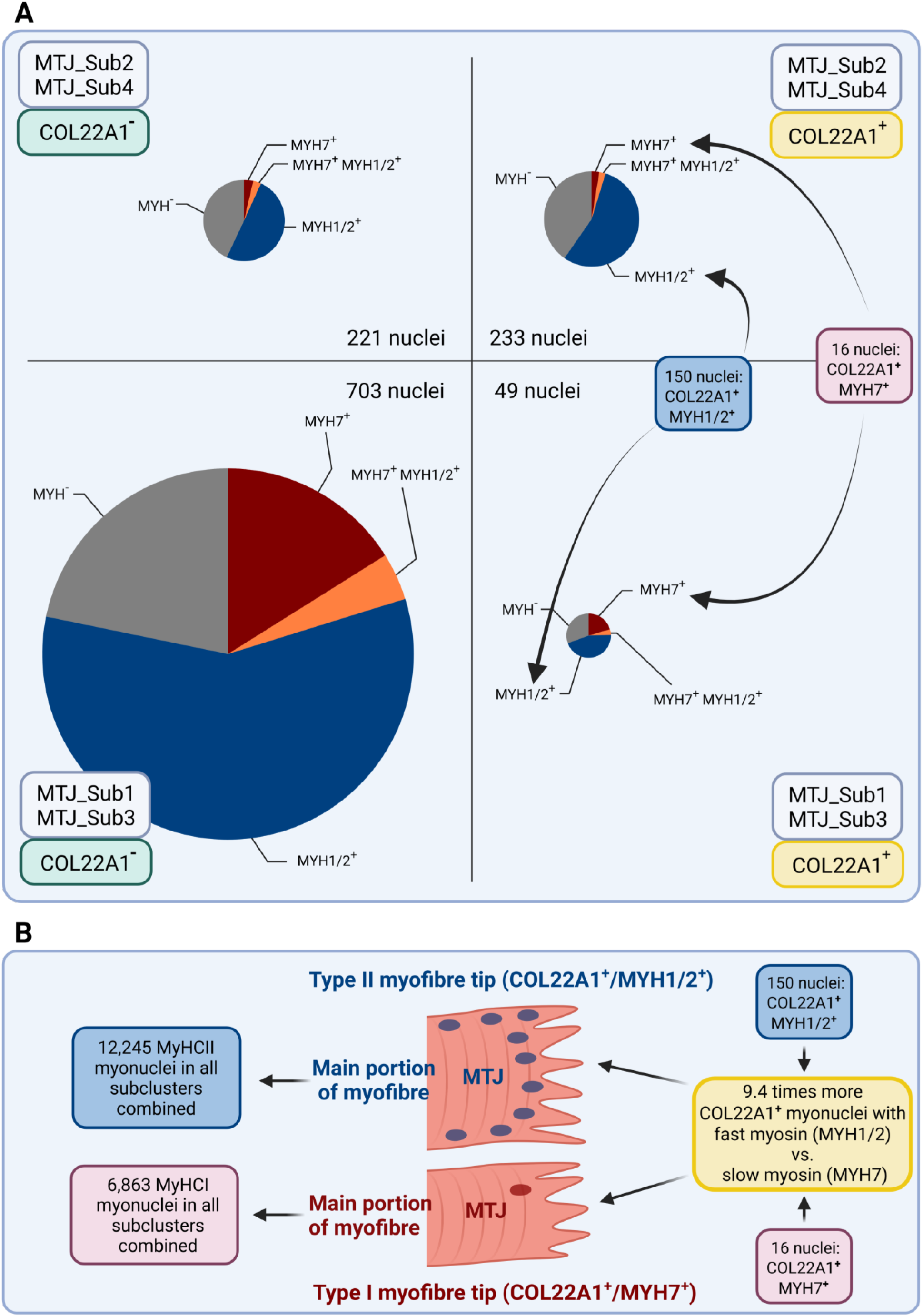
**A:** Schematic illustration of the expression of *COL22A1, MYH1, MYH2* and MYH7 in MTJ-myonuclei subclusters. The 4 MTJ-myonuclei subclusters were pooled in pairs based on either high (MTJ_Sub2/MTJ_Sub4, upper quadrant) or low (MTJ_Sub1/MTJ_Sub3, lower quadrant) expression of *COL22A1*. All *COL22A1*^+^ nuclei are accounted for in the quadrants to the right, and all COL22A1 nuclei in the quadrants to the left. Note the low abundance of *COL22A1*^+^ MTJ-myonuclei with MYH7-expression (highlighted in the two quadrants on the right). The raw-data can be found in Supplementary Data S4. **B:** Illustration of the disproportional prevalence of *MYH7*^+^ vs *MYH1/2*^+^ MTJ-myonuclei with *COL22A1* expression. Note the numbers in this figure are taken from the RAW data, and application of appropriate correction factors resulted in a 4.2 times higher prevalence (see methods for details). Elements of this figure were created with BioRender.com.

### Spatial transcriptomics of human muscle-tendon tissue

Spatial transcriptomics was performed on cryosections from 4 individuals, where each pre-defined area contained a cross section and longitudinal section from the same sample. The high level of autofluorescence of tendon tissue was a useful guide for determining the position of muscle and tendon on the sections (Supplementary Figure S8A). Together, in all 4 samples, we identified 33,215 features across 6,599 unique capture spots (Supplementary Figure S8B). Key muscle (*MYH7*) and tendon (*COMP*) transcripts were distributed clearly according to their respective tissues, while the position of *COL22A1*, the MTJ marker, was unclear due to a low number of reads (Supplementary Figure S8C). Six clusters were detected (Supplementary Figure S8D and S8E). Differentially expressed genes were only found for clusters 2 and 4, which were characterized by genes for muscle (*ACTA1, TTN, TPT1, GAPDH*) and tendon (*COMP, DCN, PRELP, THBS4*), respectively (Supplementary Data S5).

### Mapping snRNA-seq subclusters onto spatial transcriptomics data

The four snRNA-seq MTJ-myonuclei subclusters corresponded with the spatial transcriptomics rendering of the muscle-tendon tissue interface (Figure 5B, Supplementary Figure S9). Furthermore, FibTen_Sub5 (and to a lesser extent FibTen_Sub3) clearly mapped to the tendon proper, which unequivocally identifies FibTen_Sub5 as a population of pure tenocyte nuclei (Figure 5B, Supplementary Figure S9). This is in line with the lack of concordance between FibTen_Sub5 with previously reported cell clusters derived from pure skeletal muscle (Supplementary Figure S5A-B), while appearing in human scRNA-seq data from muscle/tendon/MTJ (Supplementary Fig S5B, see Yan et al. 2022) and human scRNA-seq data from pure tendon (Supplementary Fig S5B, see Kendall et al. 2020). For the remaining cell clusters (endothelial cells, smooth muscle cells, and immune cells), the spatial analysis showed a ubiquitous distribution across muscle and tendon tissue (Supplementary Figure S9).

## Discussion

More than 30 years ago, protein enrichment of talin and desmin at the MTJ was reported by James G. Tidball (Tidball et al. 1986; Tidball 1992), and over a decade later Manuel Koch and colleagues identified collagen XXII as a marker of the MTJ in skeletal and cardiac muscles (Koch et al. 2004). Unbiased discovery of new proteins at the MTJ is challenging due to difficulties in isolating this narrow anatomical region comprising two distinct tissues, and mapping of the proteins and RNAs at the MTJ is still incomplete. Here, by integrated snRNA-seq and spatial transcriptomics of healthy human muscle-tendon tissue, we identify 99 DEGs across four distinct clusters of myonuclei at the MTJ. From these genes, we demonstrate 8 MTJ-enriched proteins by immunofluorescence. The specific roles of these newly discovered proteins at the MTJ is unknown, but they cover a wide range of functions, from actin assembly and F-actin formation (ABLIM1, FHOD3), sarcomere organization (FHOD3), basement membrane adhesion (FRAS1, FREM2), intracellular cargo transport (BICD1), metallopeptidase activity (ADAMTSL1), macrophage differentiation (CPM), peptide hormone degradation at the cell surface (CPM), extracellular matrix organization (ABI3BP) and cell-substrate adhesion (ABI3BP). Based on nuclei profiles, we also identify a transitional region between the MTJ and the myofibre proper, in addition to a population of MTJ mononuclear cells.

The main focus on the MTJ-myonuclei in the present study was inspired by the consistent reporting of a specialized cluster of MTJ-myonuclei in recent snRNA-seq studies in mice (Dos Santos et al. 2020; Kim et al. 2020; Petrany et al. 2020; Wen et al. 2021). Here, we present the first transcriptional profile of human MTJ-myonuclei, constituting 47 DEGs, including transcripts for the known MTJ-enriched proteins, collagen XXII (*COL22A1*) and neural cell adhesion molecule 1 (*NCAM1*). Notably approximately half of these DEGs were not reported in the rodent MTJ-myonuclei, and only 6 DEGs were enriched in human and all the rodent MTJ-clusters (Dos Santos et al. 2020; Kim et al. 2020; Petrany et al. 2020; Wen et al. 2021), highlighting the importance of studying human tissue, and also underlining the differences that can be observed in snRNA-seq data, probably related to factors such as methods of nuclei isolation, choice of muscles and bioinformatics approach.

According to our UMAP and IF staining patterns of proteins differentially expressed by the 4 MTJ-subclusters, the *COL22A1*^*+*^ MTJ_Sub2 and MTJ_Sub4 subclusters represent a population of myonuclei belonging to the collagen XXII defined MTJ-domain. The low expression of *COL22A1* we observed in MTJ_Sub1 and MTJ_Sub3 has previously been reported in some (Kim et al. 2020) or all (Wen et al. 2021) MTJ-subclusters. However, by IF we could couple the gene expression in MTJ_Sub2/MTJ_Sub4 (*ABLIM1, COL22A1, CPM, FRAS1, FREM2, NCAM1*) with protein enrichment specifically at the MTJ, and the gene expression in MTJ_Sub1/MTJ_Sub3 (*ABLIM1, NCAM1*) with protein enrichment in the transitional myofibre region. While the presence of a transitional region of the myofibre has been indicated at the protein level by consistent IF staining of NCAM1 at the MTJ also in regions without collagen XXII (Jakobsen et al. 2018, 2021; Karlsen et al. 2022), this is to our knowledge the first evidence of a transcriptionally corresponding population of *COL22A1*^*−*^ */NCAM1*^+^ myonuclei. The high expression of *ABLIM1* in all 4 MTJ-subclusters together with the IF staining pattern of the ABLIM1 protein places ABLIM1 alongside NCAM1 in this transitional region and at the MTJ. Other DEGs in the 4 subclusters could share similar properties regarding the collagen XXII defined MTJ-domain and the transitional domain. It is worth noting that the snRNA-seq dataset appears to be validated by the IF confirmations and spatial transcriptomics, placing the MTJ clusters (and others) spatially as expected. Together these findings expand our understanding of the compartmentalization at the MTJ from gene expression to protein enrichment and raises the question whether such a transitional region also exists adjacent to the postsynaptic myonuclei at the NMJ.

In addition to the focus on the MTJ, a few observations are worth noting. Firstly, satellite cell nuclei were only found in a single well-defined cluster, indicating the lack of states or subtypes of satellite cells at the MTJ under basal conditions. Secondly, 15 of the 99 enriched genes in the MTJ-myonuclei are designated as matrisome proteins (Naba et al. 2012), which is interesting as these matrisome genes were enriched in myonuclei. In further support of a role for myonuclei in maintaining the muscle ECM, it is worth noting that *LAMA2*, encoding the muscle-specific laminin known as merosin (Patton et al. 1997), was enriched in clusters representing the MTJ-myonuclei, fibroblasts, satellite cells and tenocytes (Figure 2B, Supplementary Data S1), revealing a joint contribution of these cell types and myonuclei to basement membrane regeneration (Vracko and Benditt 1972; Mackey and Kjaer 2017a). Finally, enriched genes in type I or II myonuclei support proteomics profiles of single human myofibres (Murgia et al. 2021), including structures such as sarcoplasmic reticulum (*ATP2A2, ATP2A1*), and sarcomere (*MYOM3, TNNC2, TNNT3, TNNI2, TPM1, TPM3*) (Supplementary Data S1, see the tabs “type I” and “type II”). MYOM3 was the only sarcomeric gene we found to be enriched at the MTJ, where it was detected in the MTJ_Sub3 cluster, approaching the *MYH7* clusters on the UMAP. We find the unexpectedly low prevalence of *COL22A1^+^ /MYH7*^+^ myonuclei intriguing. Whether this can be explained by a lower turnover rate of MTJ-proteins in slow myofibres is unknown, but these data clearly suggest further specificity between fast and slow myofibres in the MTJ-domain. In relation to sarcomeric proteins, we found RBM20 enriched in all 4 MTJ subclusters (Supplementary Data S1). RBM20 is a potent regulator of titin splicing (Riley et al. 2022), which suggests structural specialization of the myofibre tips at the level of the sarcomere.

The DEG signature of each of the FibTen subclusters serves to further underline the intrinsic differences between muscle fibroblasts and tenocytes, as recently reported at the protein level in extracellular vesicles isolated from human tenocytes and muscle fibroblasts (Yeung et al. 2020). The spatial transcriptomics and cross-comparison with cell-clusters in human snRNA/scRNA-seq data clearly pointed towards FibTen_Sub5 as a pure tenocyte cluster. FibTen_Sub3 transcripts seemed to be more spatially concentrated at the periphery of the tendon (Supplementary Figure S9), and was the best match to a recently reported muscle-tendon progenitor subpopulation (Supplementary Figure S5B) (Yan et al. 2022). The FibTen_Sub3 also demonstrated a high representation of the MTJ-enriched proteins we discovered by LCMS-proteomics, of which CILP, ITGA10, and THBS4 were confirmed by IF analysis (Karlsen et al. 2022), as well as HMCN1 recently shown at the MTJ (Welcker et al. 2021), and the muscle fibre specific LAMA2 (Supplementary Data S3). FibTen_Sub3 was also the non-myonuclei cluster containing the most *COL22A1*^+^ nuclei (Supplementary Data S4). The presence of a *COL22A1*-expressing tenocyte population has previously been reported in scRNA-seq (Scott et al. 2019) and snRNA-seq in mice (Dos Santos et al. 2020; Petrany et al. 2020; Wen et al. 2021), and together our data indicate that the FibTen_Sub3 cluster represents a special population of peripheral tendon cells, or myotenocytes (Scott et al. 2019), responsible for maintenance of the tendon side of the MTJ. Together these findings on fibroblasts and tenocytes provide insight into the cells responsible for maintaining the matrix of muscle, tendon, and the specialized zone of the muscle-tendon interface.

In conclusion, through profiling the human muscle-tendon interface at the single nucleus level we advance the understanding of this specialized zone, a key region in muscle pathophysiology. Myonuclei in the MTJ cluster of dystrophin-deficient mice show many genes with abnormal gene expression pattern compared to the other myonuclei in the fibre type-specific clusters (Chemello et al. 2020), and the MTJ is resistant to experimental molecular disruption of muscle proteins (Cohn et al. 2002). Furthermore, MTJ strain injuries, induced by explosive high force movements, have poor clinical prognosis with a high rate of re-injury. Clearly, many gaps remain in our understanding of MTJ regulation and repair. To this end, our dataset provides a detailed reference for gene expression at the single nucleus level of the human MTJ, in the context of the full repertoire of mononuclear cells and myofibre types in healthy human muscle and tendon tissue.

## Supporting information

Figure 1_BioRxiv Publication License

Figure 5_Background - BioRxiv Publication License

Figure 5_Cells and proteins overlay - BioRxiv Publication License

Figure 5_Cells and proteins overlay - BioRxiv Publication License

## Acknowledgments

The authors would like to thank Anja Jokipii-Utzon and Ann-Christina Ronnié Reimann for technical assistance. We also acknowledge Irina Korshunova and BRIC’s Single-Cell Genomics Core Facility; Rajesh Somasundaram and BRIC’s Flowcytometry Core Facility; the Core Facility for Integrated Microscopy, Faculty of Health and Medical Sciences, University of Copenhagen. This work was supported by Nordea Foundation (Center for Healthy Aging Grant to M.Kj.), Novo Nordisk Foundation (NNF20OC0064829) to A.L.M., The Lundbeck Foundation (R303-2018-3427) to A.K., The Lundbeck Foundation (R344-2020-254) to A.L.M., and by the Deutsche Forschungsgemeinschaft (FOR2722) to M.Ko.

The authors declare no conflicts of interest, financial or otherwise.

## Author contributions

A.L.M, S.S., A.K., C.C.Y., P.S. and M.Kj. conceived the idea and contributed to conceptual design. M.R.K., J.R.J. and M.Ko. provided resources. A.K., C.C.Y. and P.S. developed and performed the snRNA-seq. L.D. and P.S. developed and performed the spatial transcriptomics. A.K., C.H. and A.L.M. developed and performed the IF. A.K., P.S., S.S. and A.L.M. performed data interpretation and wrote the manuscript. All authors edited and approved the manuscript.

## Methods

### EXPERIMENTAL MODEL AND SUBJECT DETAILS

#### Human subjects and sample collection

Semitendinosus muscle-tendon tissue from healthy patients was collected for snRNA-seq from three male patients, for spatial transcriptomics from 2 male and 2 female patients, and gracilis muscle-tendon tissue for single fibre microscopy from 1 male and 1 female patient (age 20-44 years, height 165-185 cm, weight 66-97 kg, BMI 20.9-29.0, non-smokers with no known disease such as diabetes, arthritis or blood borne diseases), all scheduled for anterior cruciate ligament reconstruction surgery with hamstring tendon grafts. None of the patients had performed regular heavy resistance exercise with the hamstring muscles in the months prior to the surgery. The study was approved by the Research Ethics Committees of the Capital Region of Denmark (ref. H-3-2010-070 and ref. H-20044907) and conformed to the Declaration of Helsinki II. All patients gave written informed consent before inclusion to the study.

On the day of the surgery, excess semitendinosus tissue was collected within 8 minutes in a precooled 50 ml falcon tube with 1% ice cooled PBS, and was kept on ice while being divided into smaller pieces (∼50 mg, Figure 1). Tissue for snRNA-seq was transferred to cryotubes and immediately frozen in liquid nitrogen within maximum 40 minutes from the time of extraction from the body before being stored at −80 °C. All the samples contained a piece of tendon with the muscle fibres attached. Tissue for spatial transcriptomics and immunofluorescence was carefully aligned and embedded in Tissue-Tek, and frozen in isopentane pre-cooled in liquid nitrogen before storage at −80 °C. Gracilis single fibres for immunofluorescence were fixed as described (Ralston et al. 1999; Mackey and Kjaer 2017b).

### METHOD DETAILS

#### Nuclei isolation

The samples were removed from the −80 °C freezer and transferred on dry-ice to a cryochamber (−20 °C) where a total of approximately 500 mg tissue was cut into fine pieces with a scalpel (Figure 1A), before being divided evenly into ten 2 ml screwcap tubes (522-S, Techtum, Nacka, Sweden), each containing five 2.3 mm stainless steel balls (BioSpec Products, Bartlesville, Oklahoma, USA), and stored at −80 °C.

The nuclei are isolated using the Nuclei EZ Prep kit (NUC101, Sigma-Aldrich, Saint Louis, Missouri, USA), as follows. 1 ml Nucleic EZ lysis buffer was added to each tissue tube and immediately shaken for 20 seconds at 4 m/s in a FastPrep-24 (MP Biomedicals, Illkirch, France), followed by 5 min cooling on ice. The shaking and cooling were repeated and the homogenate was transferred to a new tube and gently mixed with 0.5 ml Nuclei EZ lysis buffer, using a 1000 μl pipette. The tubes were incubated for an additional 5 min on ice with two gentle mixings. Then the homogenate from all ten tubes was filtered through a 70 μm mini strainer (PluriStrainer Mini, pluriSelect Life Science, Leipzig, Germany), including a wash with 1.5 ml Nuclei EZ lysis buffer. The nuclei were spun down (500 g, 5 min, 4°) and the pellet resuspended in 1.5 ml Nuclei EZ lysis buffer by mixing 10 times with a 1000 μl pipette. The nuclei were spun down (500 g, 5 min, 4 °C), 0.5 ml nuclei wash, and suspension buffer (NWS-Buffer: 2% BSA, 2 mM MgCl_2_, 0.5% Protector RNAse Inhibitor (SKU3335399001, Merck, Soeborg, Denmark)) was carefully added, followed by 5 min incubation on ice before resuspending the nuclei in 1.0 ml NWS-Buffer. The wash was repeated, and the nuclei resuspended in 1 ml NWS-buffer, followed by filtering through a 40 μm mini strainer (PluriSelect Mini), including a wash with 0.5 ml NWS-buffer. The nuclei were pelleted again and resuspended in 1.5 ml NWS-buffer. 1.2 ml was pelleted and resuspended in 300 μl NWS-buffer. The nuclei were stained with 7-AAD and sorted by FANS for single nuclei.

#### Single nuclei RNA sequencing

∼15,000 nuclei per sample were used for generating a snRNA sequencing library using the Chromium Next GEM Single Cell 3’ Kit v3.1 (PN-1000269, 10x Genomics, Pleasanton, CA, USA) according to the manufacturer’s protocol.

The snRNA sequencing libraries were sequenced on an Illumina NovaSeq 6000 machine, ∼450M read pairs with 2×150 bp sequencing by GENEWIZ (Leipzig, Germany).

One sample failed the cDNA synthesis and therefore the final sequencing data represent single nuclei from three individuals.

#### Analysis of single nuclei RNA sequence data

The read sequences were paired and quality filtered using FastP v0.19.4 (Chen et al. 2018) with default settings (adaptor trimming disabled), followed by alignment to the transcriptome and read counting using Cell Ranger count v3.1.0 (10xGenomics). The transcriptome was generated from the human genome assembly GRCh38 with introns included in the transcripts to obtain pre-mRNA sequences.

Identification of nuclei containing droplets and reduction of background from ambient RNA was achieved using CellBender v0.2.0 (Fleming et al. 2022) with recommended settings. Nuclei doublets within the droplets were removed by using Scrublet vDec2020 (Wolock et al. 2019); three rounds as described for tissue. The metrics are shown in Table 1.

The three samples were combined using the Seurat v4.1.0 package (Stuart et al. 2019) by first normalizing each dataset with SCTransform and then integrate the data using FindIntegrationAnchors and IntegrateData. Main nuclei types were defined using FindNeighbours and FindClusters based on PCA (30 dimensions). Subtypes were found by repeating the clustering separately on selected main types (MTJ, MyHCI, MyHCII, Immune, Fib^+^Ten, Smooth, or Endo). Data are deposited in the ArrayExpress database accession number E-MTAB-12529.

Markers for each of the clusters (DEGs, differentially expressed genes) were found using Seurat FindConservedMarkers selecting markers conserved for all three samples (all three FC>2 and FDR < 0.05).

Markers lists for clusters in published single cell/nuclei RNA-seq experiments were used to compare with the present clusters using the Seurat AddModuleScore function.

#### Calculation of correction factors *COL22A1*^*+*^*/MYH7*^*+*^ nuclei

The observed imbalance between the 9.4 times higher proportion of *COL22A1*^+^ */MYH1/2*^+^ nuclei (150 nuclei) vs. *COL22A1^+^ /MYH7*^*+*^ nuclei (16 nuclei) was based on transcript counting at single nuclei level, which could lead to systematic biased conclusions. Therefore, we first applied a false-negative correction factor to account for the prevalence of MYH7-negative myonuclei within in the MyHCI cluster and *MYH1/2*-negative myonuclei within the MyHCII cluster.

Within the total of 6,862 myonuclei in the MyHCI clusters, 73.9% expressed MYH7. The false negative correction factor was calculated as follows: [false-negative correction factor] = [all MyCHI myonuclei] / [MyHCI myonuclei with *MYH7*-expression] = 1.35.

Within the 12,245 myonuclei in the MyHCII clusters, 93.5% expressed *MYH1* and/or *MYH2*, resulting in a false-negative correction factor for *MYH1/2* myonuclei was = 1.07.

After this, we applied a myonuclei-abundance correction factor to account for the number of MyHCI and MyHCII myonuclei in the dataset. There was 6,862 MyHCI myonuclei and 12,245 MyHCII myonuclei in the dataset. This is presumably related to the fibretype distribution and/or fibre size in the samples. To account for this, we calculated the myonuclei-abundance correction factor = [12,245 MyHCII myonuclei] / [6,862 MyHCI myonuclei] = 1.78.

The raw data were then corrected as follows:

[16 *COL22A1^+^ /MYH7*^+^ nuclei] x [MyHCI false-negative correction factor] x [myonuclei-abundance correction factor] = 16 × 1.35 × 1.78 = 38.6 COL22A1 /MYH7 nuclei.

[150 *COL22A1^+^ /MYH1/2*^+^ nuclei] x [MyHCII false-negative correction factor] = 150 × 1.07 = 160.4 *COL22A1^+^ /MYH1/2*^+^ nuclei.

After correction, the proportion of COL22A1 /MYH1/2 nuclei (160.4 nuclei in the dataset, corrected) vs. *COL22A1^+^ /MYH7*^+^ nuclei (38.6 nuclei in the dataset, corrected) was 160.4/38.6 = 4.2.

#### Spatial transcriptomics

Tendon-MTJ-Muscle sections were analyzed with Spatial Transcriptomics using the Visium Spatial Gene Expression kit (10x Genomics, Pleasanton, CA, USA). Embedded samples containing both tendon and muscle were cut in a cryostat (10 μm) parallel to the tendon as well as transverse such that each capture area contains a sample cut in both directions. The sections were stained with H&E according to the manufactures protocol and imaged for H&E (brightfield) and autofluorescence (ex470/em525) using an Axio Scan.Z1 microscope (Zeiss, Birkerød, Denmark). Tissue was digested (15 min) and mRNA converted to a Spatial Gene Expression Library as recommended by the manufacture. The libraries were sequenced (NovaSeq 2×150, ∼700 million read per library) by a commercial company (GeneWiz, Leipzig, Germany). The reads were aligned to the same transcriptome database as for the single nuclei and counted using SpaceRanger v1.3.1 (10xGenomics). The counts were analyzed with the Seurat v4.1.0 package (Stuart et al. 2019) by first lognormalizing each dataset with NormalizeData and then clustering with FindNeighbours and FindClusters based on PCA (30 dimensions). Data are deposited in the ArrayExpress database (accession number E-MTAB-12530). Cell type prediction scores were generated from the single nuclei RNA seq clusters by using the spacexr/RCTD method (default setting on log-transformed spatial data with doublet_mode=‘Full’) (version 2.0.0, Cable et al. 2022). As the Spatial transcriptomics data includes all cytoplasmic RNA, the single nuclei cluster “Cytoplasm” was excluded from the analysis as it would otherwise be the best fit for all spots. Markers for each of the clusters (DEGs, differentially expressed genes) were found using Seurat FindAllMarkers with default settings (wilcox, log2FC > 0.25).

#### Immunofluorescence and microscopy

##### Tissue cryosections

For immunofluorescence of Tissue Tek embedded semitendinosus muscle-tendon samples, 10μm thick muscle-tendon sections were cut in the longitudinal plane of the muscle fibres and tendon in a cryostat at −20°C, collected on glass slides and stored at −80°C. For staining, slides were removed from the freezer and dried at room temperature. Detailed information of antibodies, dilution, blocking and fixation is provided in Supplementary Table S2. All antibodies were diluted in 1% BSA (bovine serum albumin, IgG free) in TBS (tris-buffered saline, tris-base 0.05 mol/L, sodium chloride 0.154 mol/L, pH 7.4-7.6). In the cases of high background signal, a blocking step was included for 30 minutes prior to incubation with primary antibodies (5% goat serum, 5% donkey serum and 5% BSA in TBS). Fixation with 4% paraformaldehyde (5 min) or Histofix (12 min) was applied either prior to application of primary antibodies (day 1, incubated overnight), or after application of secondary antibodies (day 2, 60 min incubation) -the optimal protocol was tested for each primary antibody. The samples were washed 3 times 5 minutes in TBS between all protocol steps, and finally mounted with cover glasses and DAPI in the mounting medium (Molecular Probes ProLong Gold anti-fade reagent, cat. no. P36931). Images were captured on an Olympus BX51 microscope, controlled by the Olympus cellSens Software (www.olympus-lifescience.com), using 10x (0.3NA) or 20x (0.5NA) objectives and a 0.5x camera (Olympus DP71, Olympus Deutschland GmbH, Hamburg, Germany). Image size was 4080×3072 pixels, 1747×1315μm (2.33 pixels/μm) and 868×653μm (4.70 pixels/μm) for images taken with the 10x and 20x objective respectively. Images were viewed and cropped for presentation in ImageJ (version 1.53c; National Institute of Health; USA), and colour-blind friendly pseudo colours were applied to composite images.

##### Single muscle fibres

Adapting a previous protocol for isolating single muscle fibres (Ralston et al. 1999; Mackey and Kjaer 2017b), individual myofibres were teased from the gracilis tendon of the fixed sample in 50% glycerol/PBS and incubated in immunobuffer (IB: PBS, 50mM glycine, 0.25% BSA, 0.03% saponin, 0.05% sodium azide), with 0.1% Triton X-100 (cat. no. 9036-19-5, lot: STBJ9023, Sigma-Aldrich), in a 24-well plate. Primary antibodies, were diluted in IB (containing 0.1% Triton X-100) and added to the wells, followed by incubation overnight (two nights for ABLIM1) at room temperature. Fibres were washed and incubated with secondary antibodies, along with Hoechst 33342 (cat. no. H1399, Invitrogen, 1μg/ml: 1:100 dilution) for 2 h (4 h for ABLIM1). Fibres were washed and aligned in a drop of mounting medium (Molecular Probes Prolong Gold mounting, cat. no. P36930, Invitrogen) on a microscope slide, cover-slipped, and stored at −20 °C. Confocal images were acquired with a Zeiss LSM710 (Software: Zeiss Zen Black 2012) using an EC Plan-Neofluar x40/1.3NA Oil DIC M27 objective, image size 4336×4336 pixels / 922×922μm (4.70 pixels/μm). Hoechst, Alexa Fluor 488, and Alexa Flour 594 were excited by a 405-nm diode laser (30mW), a 458-nm argon laser (25mW), and a 561-nm solid-state laser (2mW), respectively. Images were viewed and cropped for presentation in ImageJ (version 1.53c; National Institute of Health; USA), and colour-blind friendly pseudo colours were applied to composite images.

## Supplementary information

**Supplementary Figure S1. snRNA-seq nuclei distribution between subjects and dotplots with general markers for cell types in the main analysis**.

**A:** Percentwise distribution of nuclei between the 17 snRNA-seq clusters within each subject.

**B:** Dotplot with general markers for cell types in the 17 snRNA-seq clusters. The size of the dot shows the proportion of nuclei in a cluster that express the gene, the colour gradient of each dot illustrates the average level of gene expression within a cluster.

**Supplementary Figure S2. Greyscale IF-images used for the merged pseudo colour images in Figure 3A-D**.

This figure provides a more detailed version of the IF-images shown in Figure 3A-D by including the raw grey scale channels, allow for more detailed evaluation. IF-staining for collagen XXII (COL22) indicates the myotendinous junction. Skeletal muscle fibres are marked with an “M” and tendon with a “T”. DAPI or Hoechst stained the nuclei. All scale bars are 50μm.

**A-D:** Show the protein enrichment of FRAS1 **(A)**, FREM2 **(B)**, BICD1 **(C)**, FHOD3 **(D)** at the tip of the muscle fibres in close association with the dystrophin (DYST) stained sarcolemma and the MTJ-marker collagen XXII (COL22). The second panel row is a higher magnification of the dotted box region in the first panel.

**Supplementary Figure S3. Greyscale IF-images used for the merged pseudo colour images in Figure 3E-H**.

This figure provides a more detailed version of the IF-images shown in Figure 3E-H by including the raw gray scale channels to allow for more detailed evaluation. IF-staining for collagen XXII (COL22) indicates the myotendinous junction. Skeletal muscle fibres are marked with an “M” and tendon with a “T”. DAPI or Hoechst stained the nuclei. All scale bars are 50μm.

**A-B:** Show the protein enrichment of CPM **(A)** and ADAMTSL1 **(B)** at the tip of the muscle fibres in close association with the dystrophin (DYST) stained sarcolemma or collagen IV (COL4) stained basement membrane and the MTJ-marker collagen XXII (COL22). The second panel row is a higher magnification of the dotted box region in the first panel.

**C:** Upper panel show ABI3BP signal is strongest in the MTJ-region on the extracellular matrix side of the muscle fibres and in the tendon. Lower panel show the clear extracellular localization of ABLIM1 at the tip of the muscle fibre in an isolated single fibre, dystrophin (DYST) stained the sarcolemma and Hoechst stained the nuclei.

**D:** Upper panel show ABLIM1 enrichment in the myofibre cytoplasm on tissue cross-section, strongest near the MTJ and gradually fading along the length of the muscle fibre. Lower panel show the same pattern in a single fibre, dystrophin (DYST) stained the sarcolemma and Hoechst stained the nuclei.

**Supplementary Figure S4. snRNA-seq nuclei distribution between subjects and dotplots with general markers for cell types in the subclusters**.

**A:** Percentwise distribution of nuclei between the 30 snRNA-seq subclusters within each subject.

**B:** Dotplot with general markers for cell types in the 30 snRNA-seq subclusters. The size of the dot illustrates the proportion of nuclei in a cluster that express the corresponding gene, the color gradient of each dot illustrates the average level of gene expression for the corresponding gene within each cluster.

**Supplementary Figure S5. Comparison of the snRNA-seq subclusters in the present study to clusters in other single cell/nuclei RNA-seq studies of human and mouse skeletal muscle and tendon**.

Marker genes provided for scRNA/snRNA-seq clusters in previous publications in the literature were used to score similarity to the clusters in the present study using Seurat AddModuleScore.

Note that scores are generated as a comparison between the present data and each individual report, and within each heatmap the colours go from zero (blue) to highest score (red). Therefore, the scores are not comparable between the different heatmaps (publications).

**S5A:** Comparison with single nuclei RNA-seq (snRNA-seq) in human tissue.

**S5B:** Comparison with single cell RNA-seq (scRNA-seq) in human muscle tissue, human muscle/tendon tissue, human tendon tissue and mouse tendon tissue.

**S4C:** Comparison with snRNA-seq and in mouse muscle/tendon tissue.

**Supplementary Figure S6. Dotplot with the 99 DEGs found in the 4 MTJ-myonuclei subclusters**

Dotplot with the 99 DEGs (differentially expressed genes) in the 4 MTJ-myonuclei subclusters shown for all 30 subclusters. The size of the dot illustrates the proportion of nuclei in a cluster that express the corresponding gene, the colour gradient of each dot illustrates the average level of gene expression for the corresponding gene within each cluster.

**Supplementary Figure S7. Immunofluorescence of fast and slow type myosin together with collagen XXII**

Triple immunofluorescence staining for slow (MYH7) and fast (MYH1/2) type myosin together with the MTJ-marker collagen XXII (COL22). The images are widefield microscopy of 10μm thick sections of human semitendinosus muscle-tendon tissue samples cut in the longitudinally plane of the muscle fibres, and shows clear enrichment of collagen XXII in all fibre types in human skeletal muscle tissue with a mixed fibre type. The scale bars are 50 μm.

**Supplementary Figure S8. Spatial Transcriptomics**

**A:** Cryosections of human muscle-tendon tissue from 4 individuals (A1-D1), stained for haematoxylin and eosin (H&E), or presented with autofluorescence signal alone. In each image, one section is cut longitudinally to the muscle fibre direction, and the other section is cut in the cross-sectional plane.

**B:** Number of counts (UMIs) and number of different features (transcripts) per spot for each section.

**C:** Location of spots containing genes representing muscle (*MYH7*), tendon (*COMP*), and MTJ (*COL22A1*) across all sections.

**D:** Spatial transcriptomics cluster UMAP.

**E:** Location of each cluster in D on all sections.

**Supplementary Figure S9. Mapping of the snRNA-seq subclusters onto Spatial Transcriptomics data**.

Mapping of the 30 snRNA-seq subclusters onto the Spatial Transcriptomics data. The images are from the cross-sectional cut muscle-tendon tissue sample in Sample A1 (see Supplementary Figure S8). Each dot represents a 100×100μm region of the tissue section. The colour intensity of each dot shows the level of overlap between snRNA-seq gene transcripts within a cluster on the spatial transcriptomic map. In the first image (MyHCI_Sub1) the autofluorescent green colour is the tendon, market with “T” and the darker muscle regions are marked with “M”.

**Supplementary Data S1**.

snRNA-seq analysis of markers (differentially expressed genes, DEGs) in each of the 17 nuclei clusters shown in Figure 2. The first tab (info-tab) includes abbreviations used in the following tabs. The last 4 tabs include data for a pooled analysis of all myonuclei, all non-myonuclei, or type I myonuclei and type II myonuclei.

**Supplementary Data S2**.

Data analysis for the Venn-Diagram in Figure 2C.

**Supplementary Data S3**.

snRNA-seq analysis of markers (differentially expressed genes, DEGs) in each of the 30 subclusters shown in Figure 4. The first tab (info-tab) includes abbreviations used in the following tabs.

**Supplementary Data S4**.

Raw-data showing expression counts for individual nuclei of:

**Tab 1)** *ABLIM1* and *NCAM1*.

**Tab 2)** *COL22A1, MYH1, MYH2, MYH7*.

The data shows the number of nuclei where the transcript was detected, this means that for each nucleus the data shows if the transcript was detected (value=1) or not detected (value=0).

**Supplementary Data S5**.

Spatial transcriptomics cluster markers (differentially expressed genes, DEGs). The first tab (info) explains abbreviations used in the following tabs.

**Supplementary Table S1**.

Complete list of all gene names, their corresponding ensemble ID and protein names.

**Supplementary Table S2**.

Complete overview of antibodies used for immunofluorescence, concentrations, blocking steps and method of fixation for all immunofluorescence staining protocols.

