## Supplemental figures for "Distinct myofibre domains of the human myotendinous junction revealed by single nucleus RNA-seq"

### Supplementary Figure S1

snRNA-seq
Percentage nuclei for each sample in the 17 clusters

A

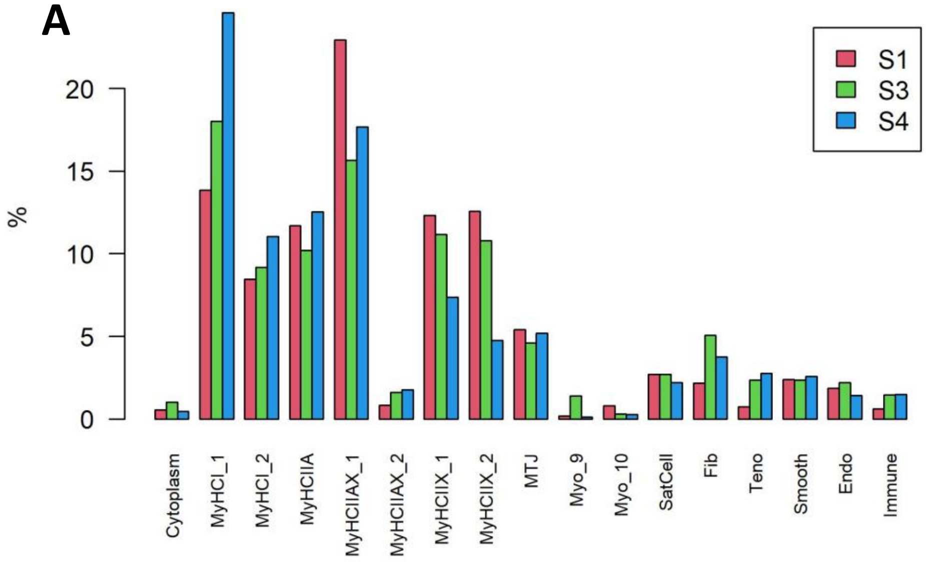


B

snRNA-seq
DotPlot with general markers for cell types in the 17 clusters

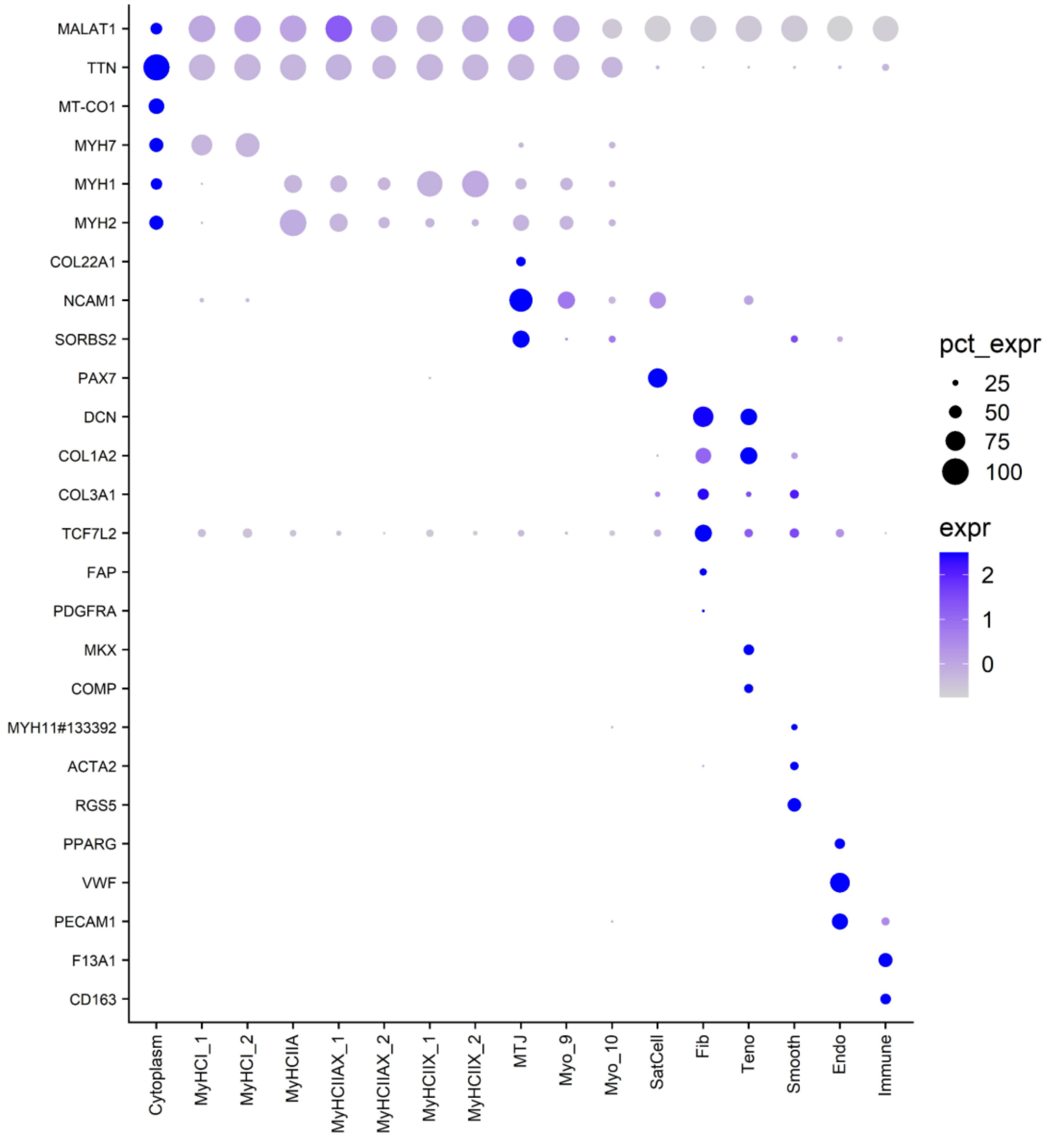

### Supplementary Figure S2

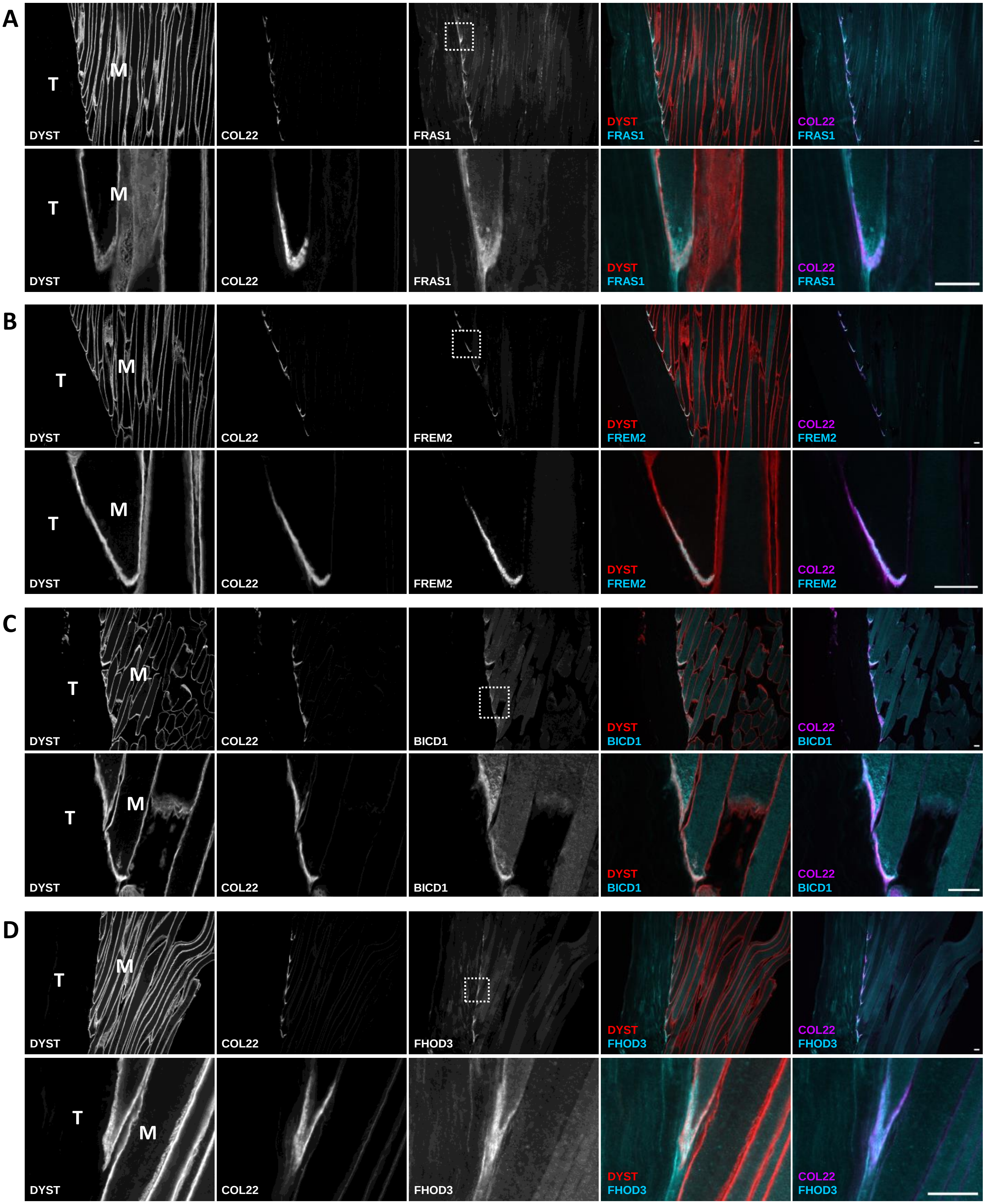

### Supplementary Figure S3

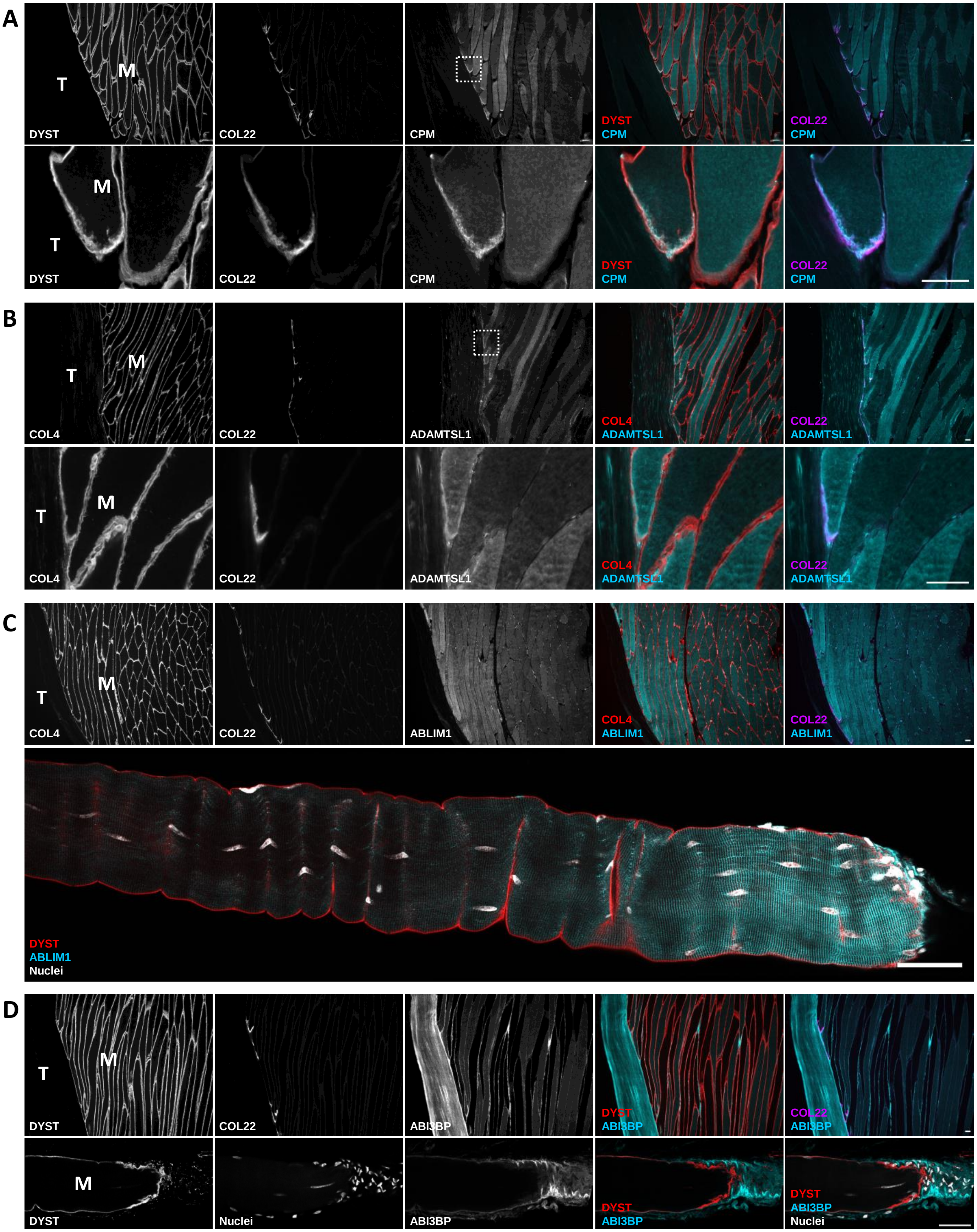

#### Supplementary Figure S4

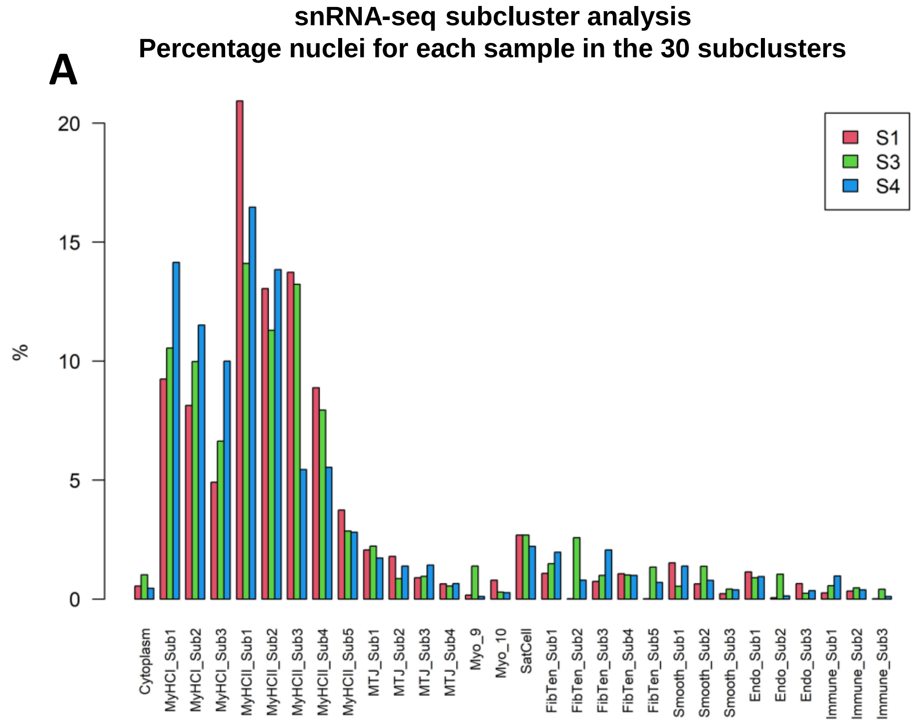


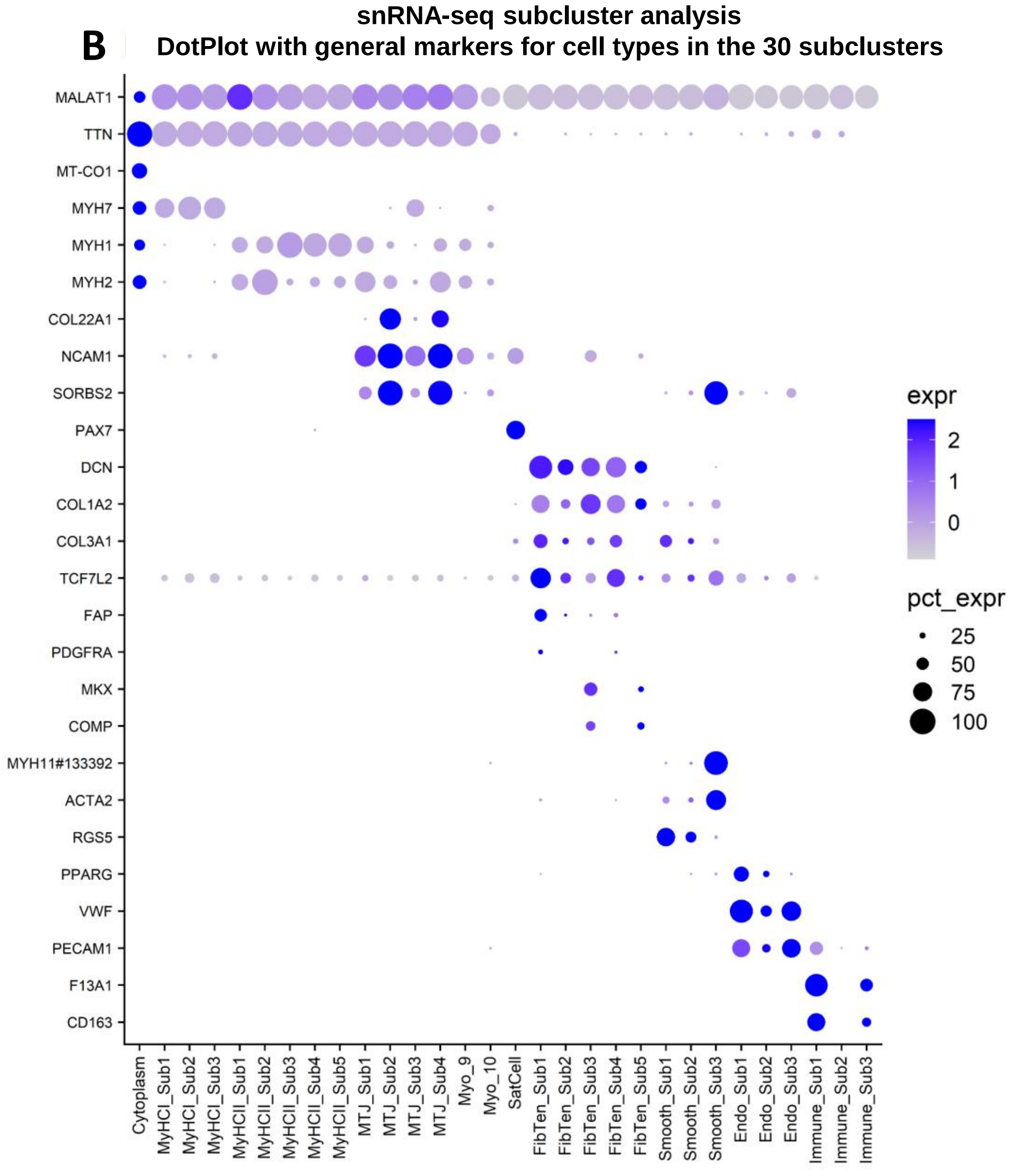

### Supplementary Figure S5A

#### Human single nuclei RNA-seq in muscle and muscle/tendon

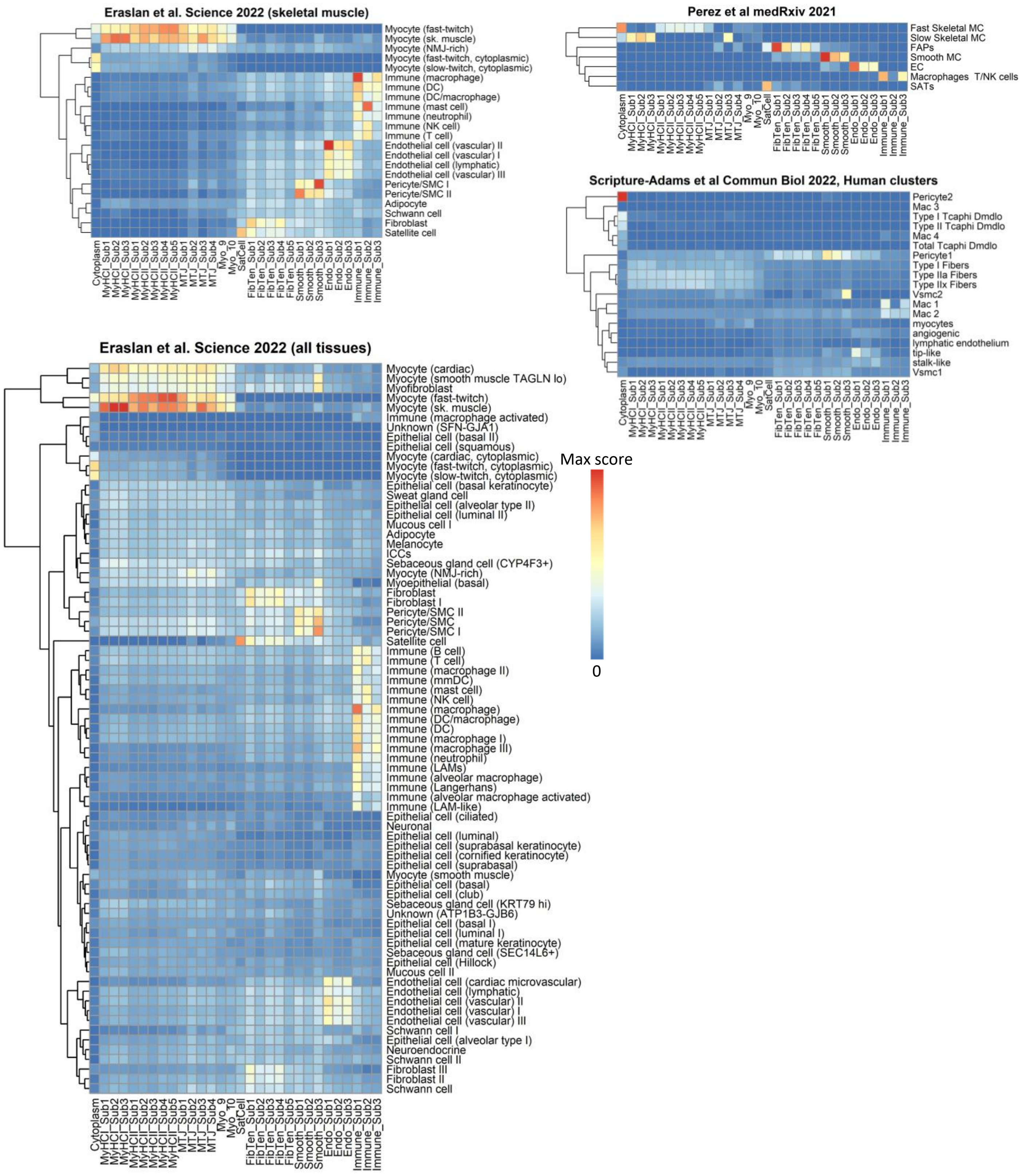

**Supplementary Figure S5B**

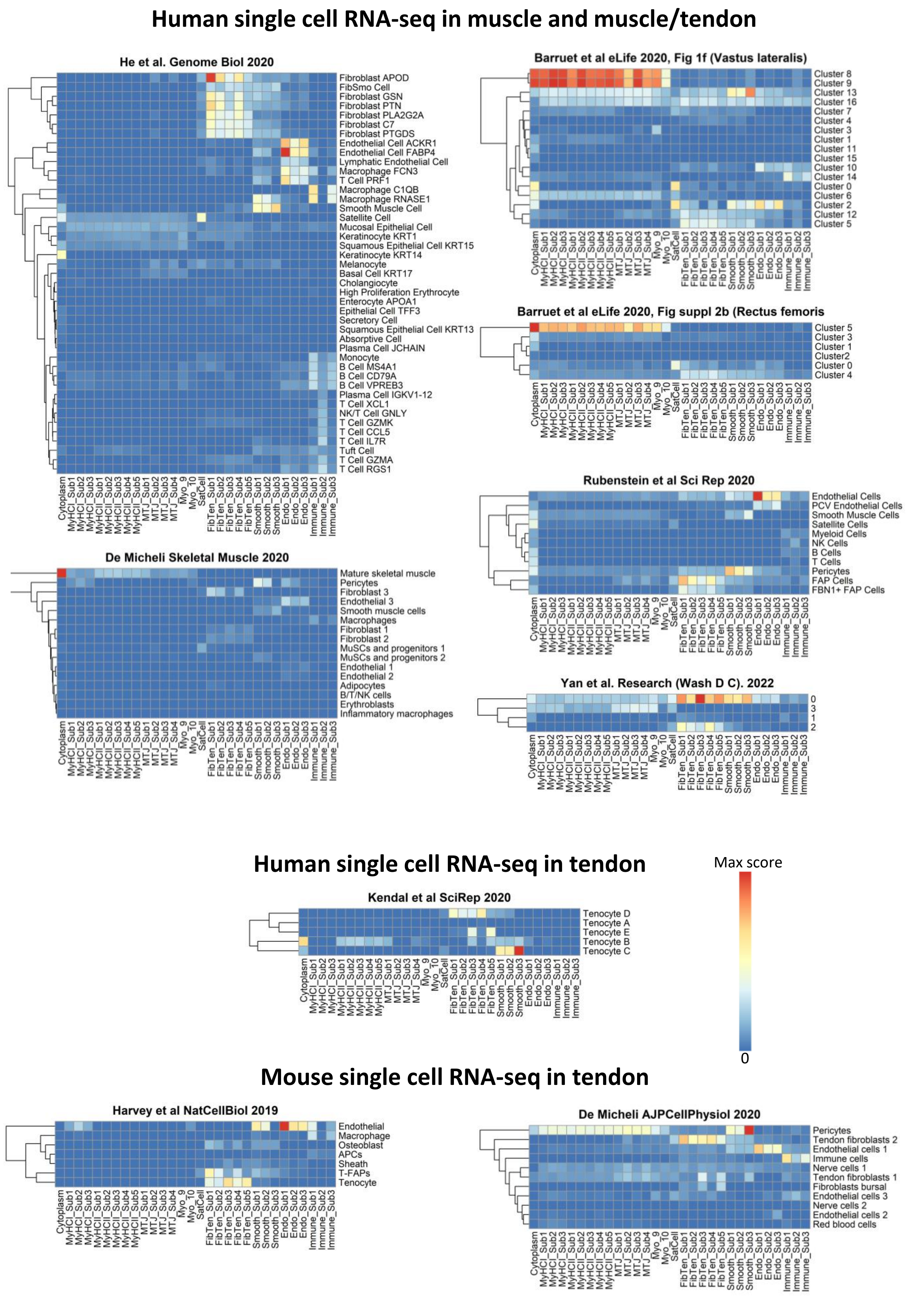

### Supplementary Figure S5C

#### Mouse single nuclei RNA-seq in muscle/tendon

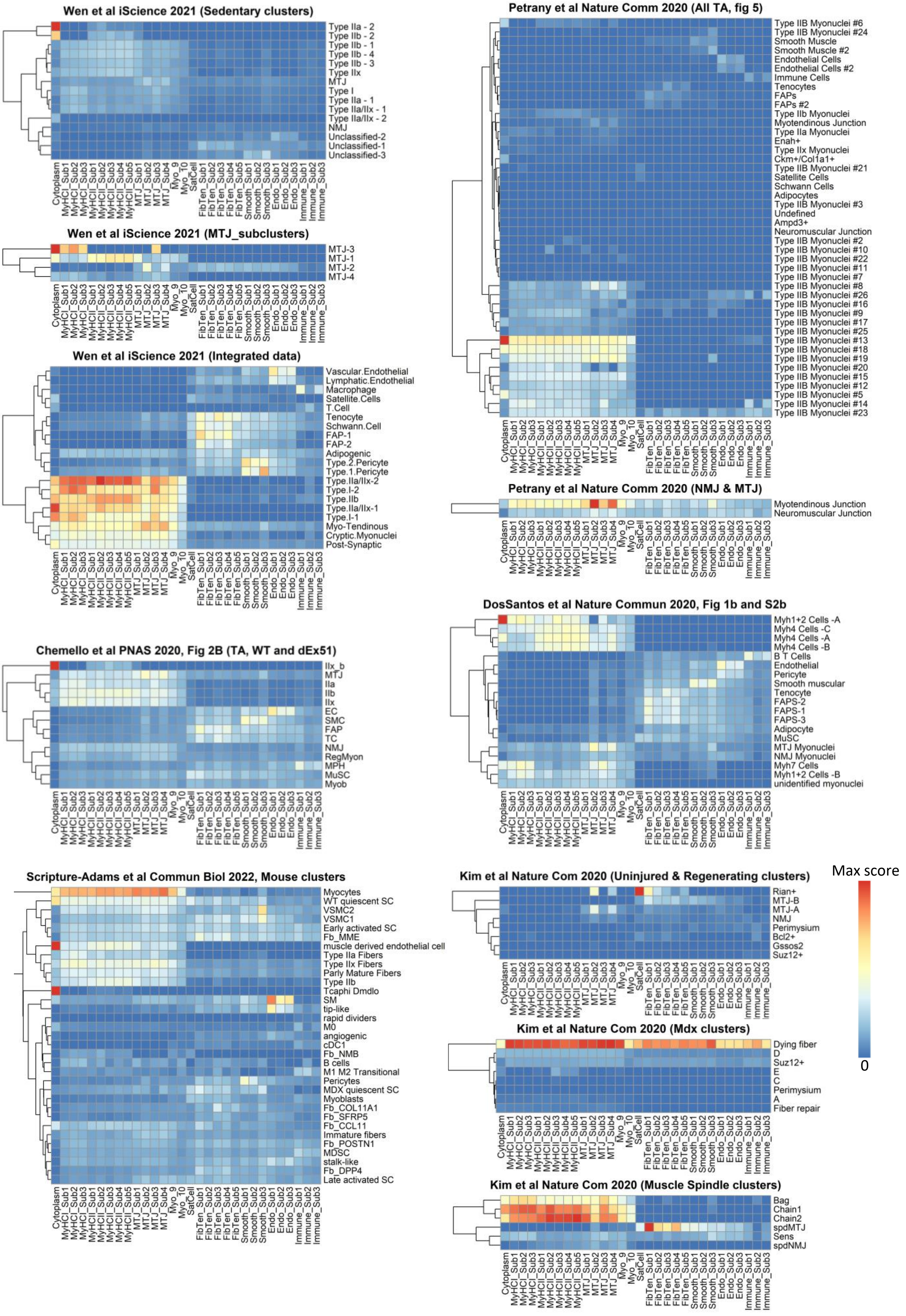

### Supplementary Figure S6

**snRNA-seq subcluster analysis**
**DotPlot with the 99 DEGs found in the 4 MTJ-myonuclei subclusters**

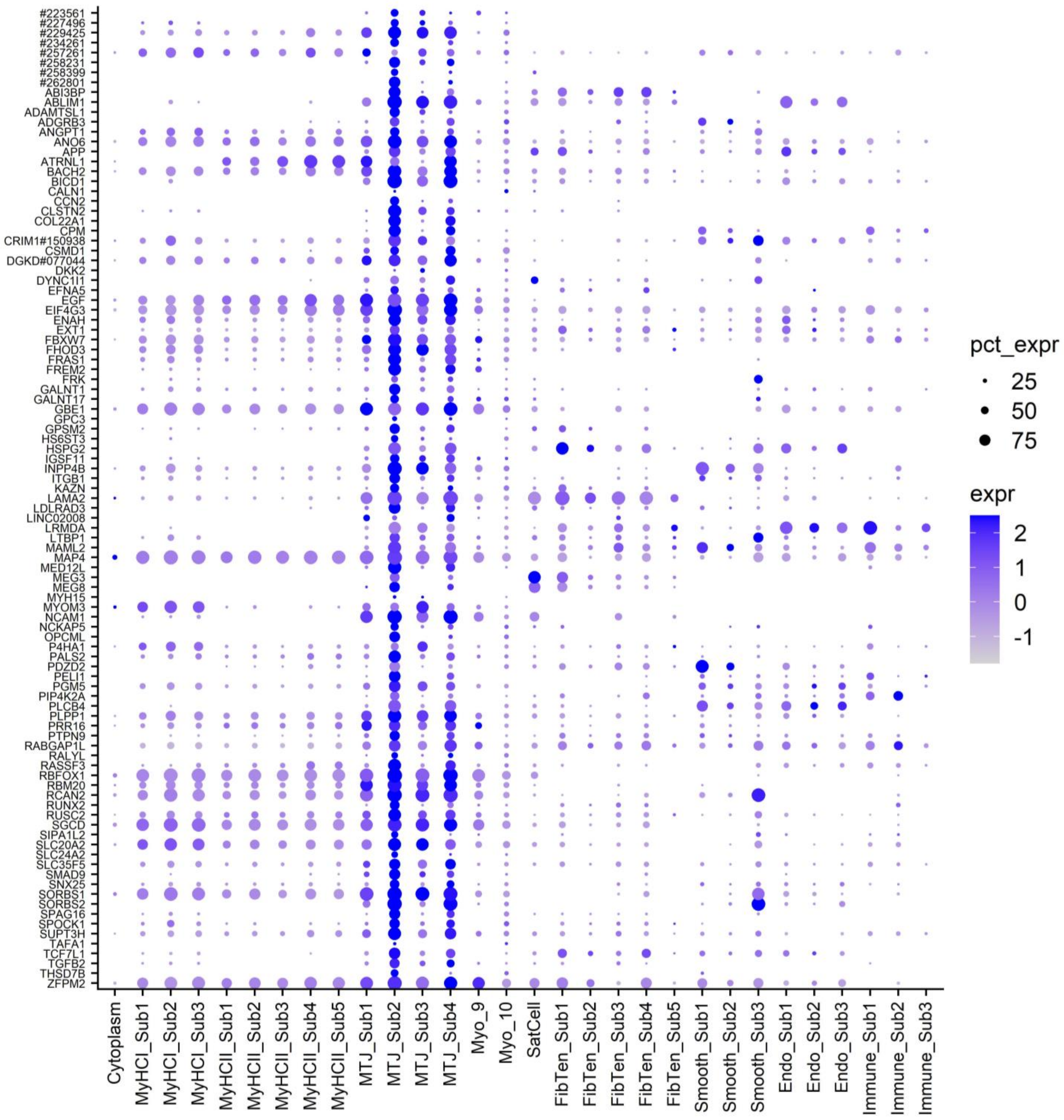

#### Supplementary Figure S7

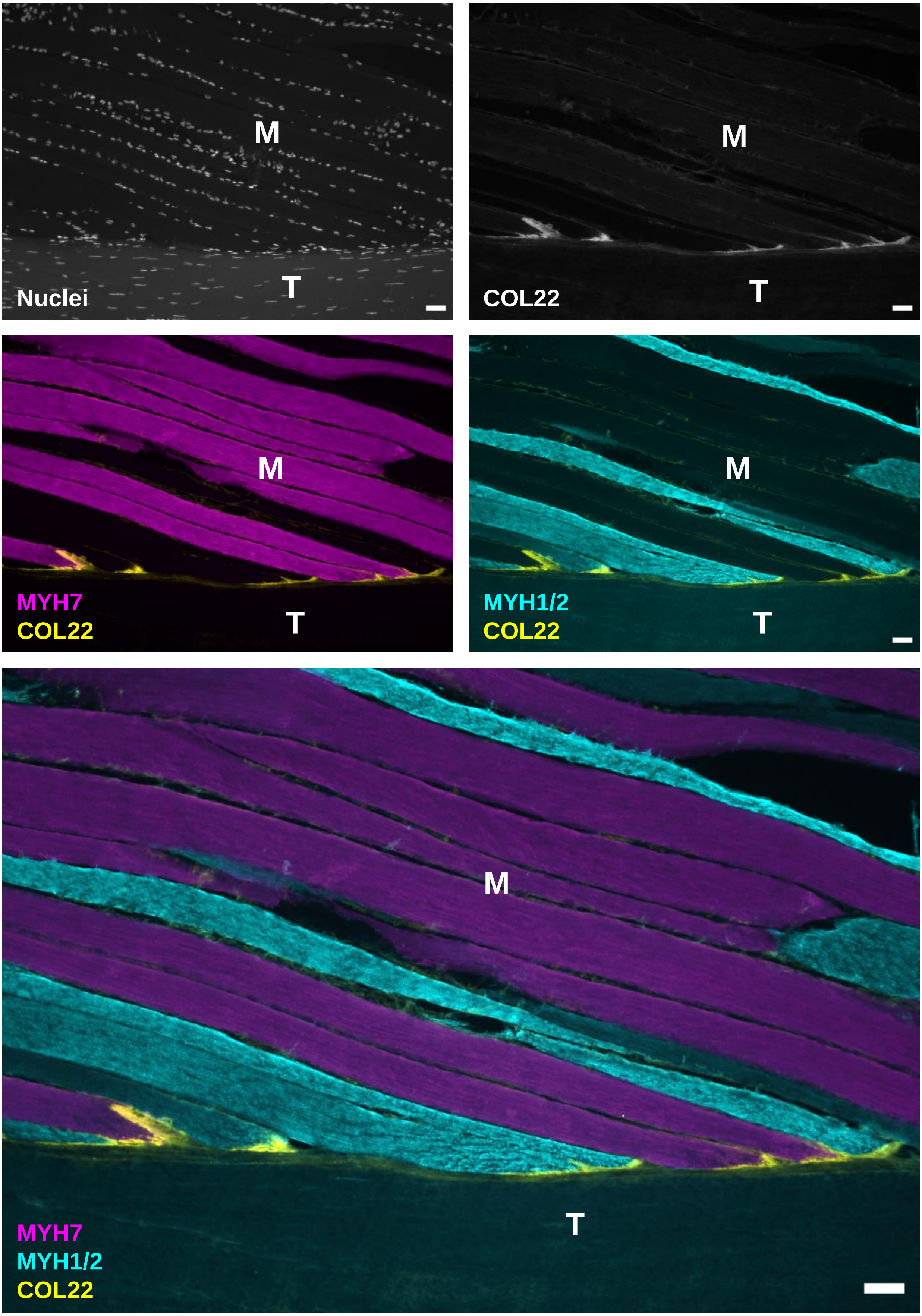

### Supplementary Figure S8

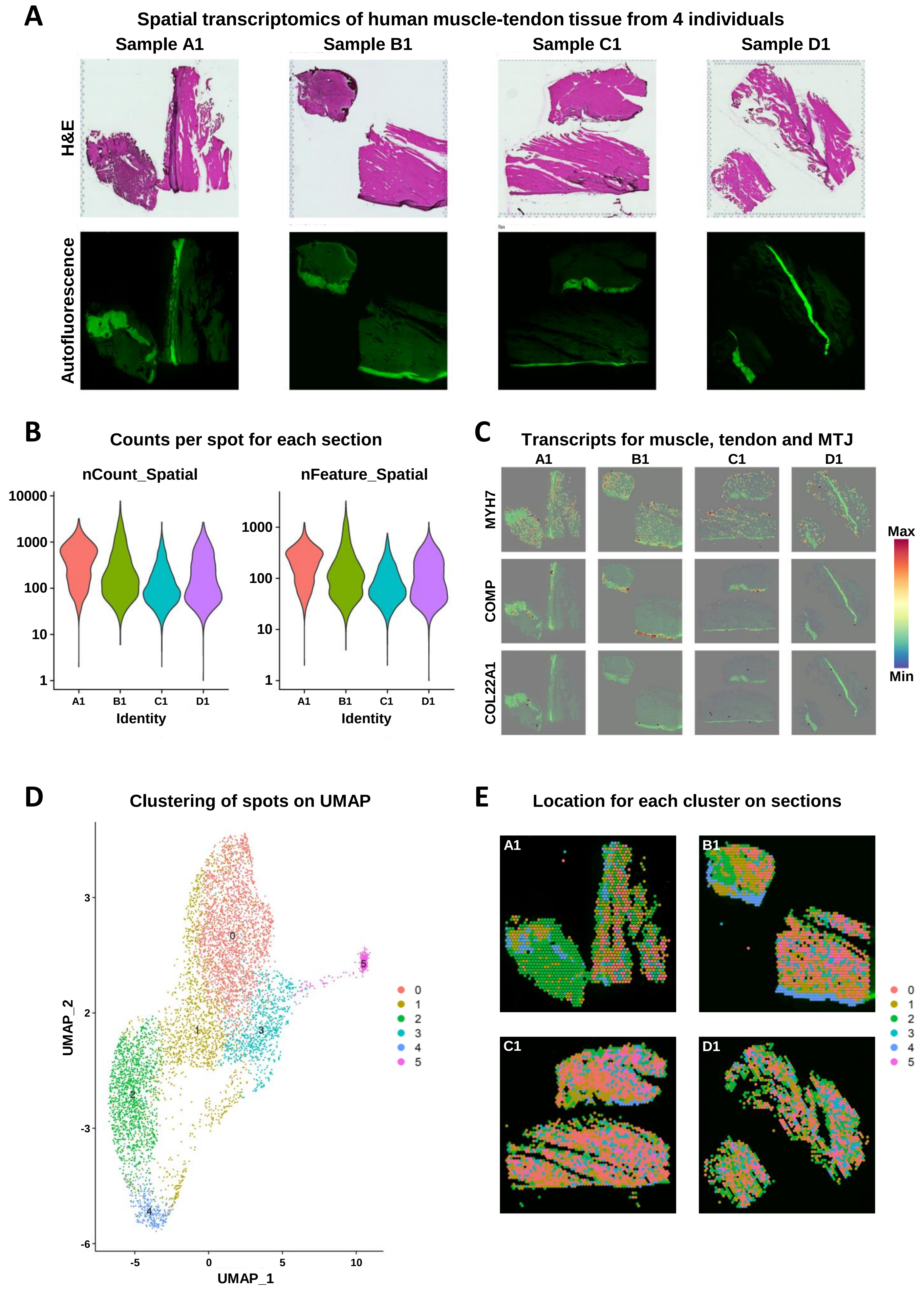

#### Supplementary Figure S9

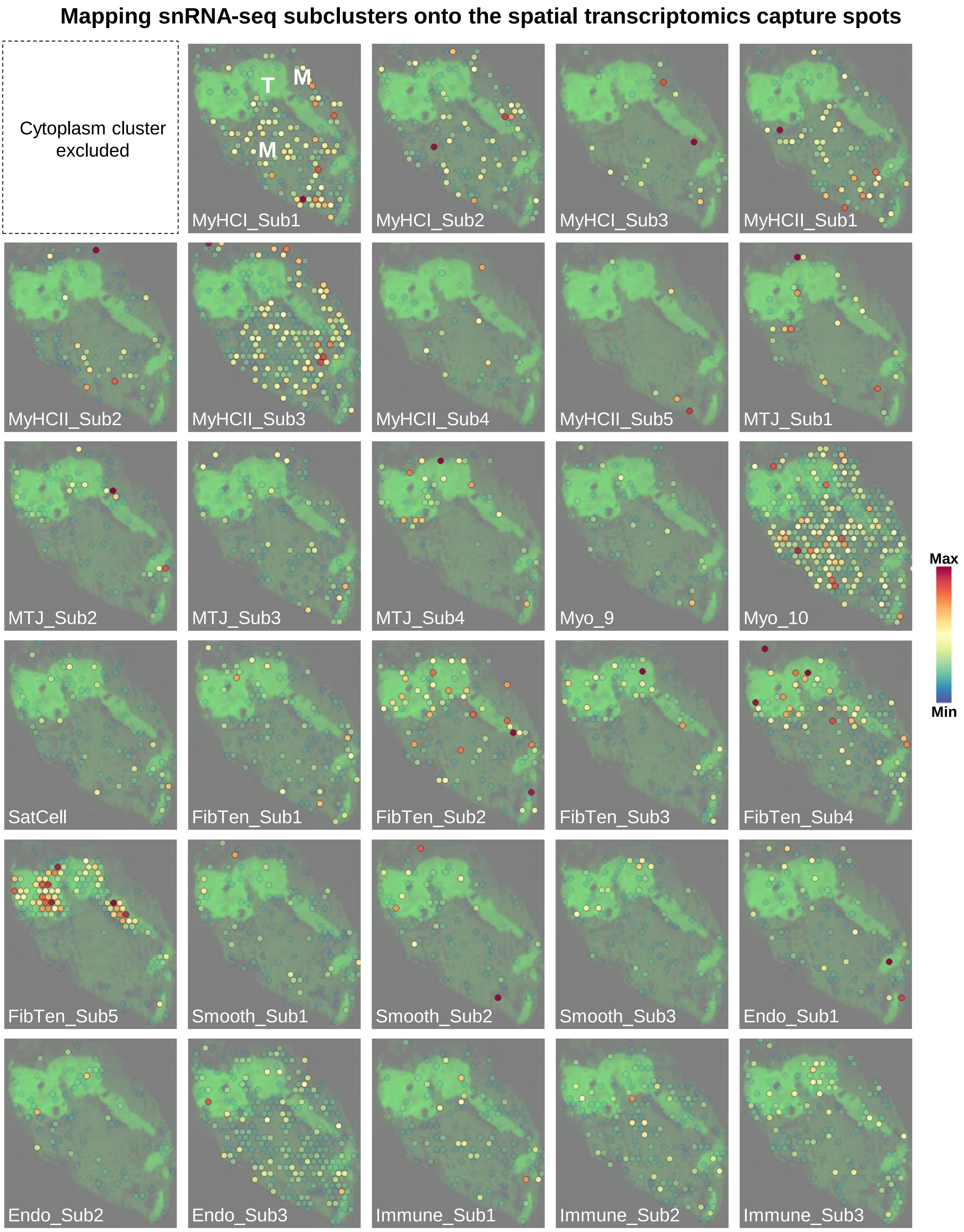
