## Supplementary Table S1 for "Distinct myofibre domains of the human myotendinous junction revealed by single nucleus RNA-seq"

**Supplemental Table S1** - Overview of gene names

| **IDName** | **Ensembl gene ID** | **Gene Name** |
| --- | --- | --- |
| *#223561* | ENSG00000223561 | N/A* |
| *#227496* | ENSG00000227496 | N/A* |
| *#229425* | ENSG00000229425 | N/A* |
| *#234261* | ENSG00000234261 | N/A* |
| *#257261* | ENSG00000257261 | N/A* |
| *#258231* | ENSG00000258231 | N/A* |
| *#258399* | ENSG00000258399 | N/A* |
| *#262801* | ENSG00000262801 | N/A* |
| *ABI3BP* | ENSG00000154175 | Target of Nesh-SH3 |
| *ABLIM1* | ENSG00000099204 | Actin-binding LIM protein 1 |
| *ACTA2* | ENSG00000107796 | Actin, aortic smooth muscle |
| *ADAMTSL1* | ENSG00000178031 | ADAMTS-like protein 1 |
| *ADGRB3* | ENSG00000135298 | Adhesion G protein-coupled receptor B3 |
| *ANGPT1* | ENSG00000154188 | Angiopoietin-1 |
| *ANO6* | ENSG00000177119 | Anoctamin-6 |
| *APP* | ENSG00000142192 | Amyloid-beta precursor protein |
| *ATP2A1* | ENSG00000196296 | Sarcoplasmic/endoplasmic reticulum calcium ATPase 1 |
| *ATP2A2* | ENSG00000174437 | Sarcoplasmic/endoplasmic reticulum calcium ATPase 2 |
| *ATRNL1* | ENSG00000107518 | Attractin-like protein 1 |
| *BACH2* | ENSG00000112182 | Transcription regulator protein BACH2 |
| *BICD1* | ENSG00000151746 | Protein bicaudal D homolog 1 |
| *CALN1* | ENSG00000183166 | Calcium-binding protein 8 |
| *CCN2* | ENSG00000118523 | CCN family member 2 |
| *CD163* | ENSG00000177575 | Scavenger receptor cysteine-rich type 1 protein M130 |
| *CHRNA1* | ENSG00000138435 | Acetylcholine receptor subunit alpha |
| *CILP* | ENSG00000138615 | Cartilage intermediate layer protein 1 |
| *CLSTN2* | ENSG00000158258 | Calsyntenin-2 |
| *COL1A2* | ENSG00000164692 | Collagen alpha-2(I) chain |
| *COL22A1* | ENSG00000169436 | Collagen alpha-1(XXII) chain |
| *COL3A1* | ENSG00000168542 | Collagen alpha-1(III) chain |
| *COMP* | ENSG00000105664 | Cartilage oligomeric matrix protein |
| *CPM* | ENSG00000135678 | Carboxypeptidase M |
| *CRIM1#150938* | ENSG00000150938 | Cysteine-rich motor neuron 1 protein |
| *CSMD1* | ENSG00000183117 | CUB and sushi domain-containing protein 1 |
| *DCN* | ENSG00000011465 | Decorin |
| *DGKD#077044* | ENSG00000077044 | Diacylglycerol kinase delta |
| *DKK2* | ENSG00000155011 | Dickkopf-related protein 2 |
| *DYNC1I1* | ENSG00000158560 | Cytoplasmic dynein 1 intermediate chain 1 |
| *EFNA5* | ENSG00000184349 | Ephrin-A5 |
| *EGF* | ENSG00000138798 | Pro-epidermal growth factor |
| *EIF4G3* | ENSG00000075151 | Eukaryotic translation initiation factor 4 gamma 3 |
| *ENAH* | ENSG00000154380 | Protein enabled homolog |
| *EXT1* | ENSG00000182197 | Exostosin-1 |
| *F13A1* | ENSG00000124491 | Coagulation factor XIII A chain |
| *FAP* | ENSG00000078098 | Prolyl endopeptidase FAP |
| *FBXW7* | ENSG00000109670 | F-box/WD repeat-containing protein 7 |
| *FHOD3* | ENSG00000134775 | FH1/FH2 domain-containing protein 3 |
| *FRAS1* | ENSG00000138759 | Extracellular matrix organizing protein FRAS1 |
| *FREM2* | ENSG00000150893 | FRAS1-related extracellular matrix protein 2 |
| *FRK* | ENSG00000111816 | Tyrosine-protein kinase FRK |
| *GALNT1* | ENSG00000141429 | Polypeptide N-acetylgalactosaminyltransferase 1 |
| *GALNT17* | ENSG00000185274 | Polypeptide N-acetylgalactosaminyltransferase 17 |
| *GBE1* | ENSG00000114480 | 1,4-alpha-glucan-branching enzyme |
| *GPC3* | ENSG00000147257 | Glypican-3 |
| *GPSM2* | ENSG00000121957 | G-protein-signaling modulator 2 |
| *HMCN1* | ENSG00000143341 | Hemicentin-1 |
| *HS6ST3* | ENSG00000185352 | Heparan-sulfate 6-O-sulfotransferase 3 |
| *HSPG2* | ENSG00000142798 | Basement membrane-specific heparan sulfate proteoglycan core protein |
| *IGSF11* | ENSG00000144847 | Immunoglobulin superfamily member 11 |
| *INPP4B* | ENSG00000109452 | Inositol polyphosphate 4-phosphatase type II |
| *ITGA10* | ENSG00000143127 | Integrin alpha-10 |
| *ITGB1* | ENSG00000150093 | Integrin beta-1 |
| *KAZN* | ENSG00000189337 | Kazrin |
| *LAMA2* | ENSG00000196569 | Laminin subunit alpha-2 |
| *LDLRAD3* | ENSG00000179241 | Low-density lipoprotein receptor class A domain-containing protein 3 |
| *LINC02008* | ENSG00000239440 | Long Intergenic Non-Protein Coding RNA 2008 |
| *LRMDA* | ENSG00000148655 | Leucine-rich melanocyte differentiation-associated protein |
| *LTBP1* | ENSG00000049323 | Latent-transforming growth factor beta-binding protein 1 |
| **IDName** | **Ensembl gene ID** | **Gene Name** |
| *MALAT1* | ENSG00000251562 | Metastasis associated lung adenocarcinoma transcript 1 |
| *MAML2* | ENSG00000184384 | Mastermind-like protein 2 |
| *MAP4* | ENSG00000047849 | Microtubule-associated protein 4 |
| *MED12L* | ENSG00000144893 | Mediator of RNA polymerase II transcription subunit 12-like protein |
| *MEG3* | ENSG00000214548 | Mediator of RNA polymerase II transcription subunit 12-like protein |
| *MEG8* | ENSG00000225746 | Maternally expressed protein 8 |
| *MKX* | ENSG00000150051 | Homeobox protein Mohawk |
| *MT-CO1* | ENSG00000198804 | Cytochrome c oxidase subunit 1 |
| *MYH1* | ENSG00000109061 | Myosin-1 |
| *MYH11* | ENSG00000133392 | Myosin-11 |
| *MYH15* | ENSG00000144821 | Myosin-15 |
| *MYH2* | ENSG00000125414 | Myosin-2 |
| *MYH7* | ENSG00000092054 | Myosin-7 |
| *MYOM3* | ENSG00000142661 | Myomesin-3 |
| *NCAM1* | ENSG00000149294 | Neural cell adhesion molecule 1 |
| *NCKAP5* | ENSG00000176771 | Nck-associated protein 5 |
| *OPCML* | ENSG00000183715 | Opioid-binding protein/cell adhesion molecule |
| *P4HA1* | ENSG00000122884 | Prolyl 4-hydroxylase subunit alpha-1 |
| *PALS2* | ENSG00000105926 | Protein PALS2 |
| *PAX7* | ENSG00000009709 | Paired box protein Pax-7 |
| *PDGFC* | ENSG00000145431 | Platelet-derived growth factor C |
| *PDGFRA* | ENSG00000134853 | Platelet-derived growth factor receptor alpha |
| *PDZD2* | ENSG00000133401 | PDZ domain-containing protein 2 |
| *PECAM1* | ENSG00000261371 | Platelet endothelial cell adhesion molecule |
| *PELI1* | ENSG00000197329 | E3 ubiquitin-protein ligase pellino homolog 1 |
| *PGM5* | ENSG00000154330 | Phosphoglucomutase-like protein 5 |
| *PIP4K2A* | ENSG00000150867 | Phosphatidylinositol 5-phosphate 4-kinase type-2 alpha |
| *PLCB4* | ENSG00000101333 | 1-phosphatidylinositol 4,5-bisphosphate phosphodiesterase beta-4 |
| *PLPP1* | ENSG00000067113 | Phospholipid phosphatase 1 |
| *PRR16* | ENSG00000184838 | Protein Largen |
| *PTPN9* | ENSG00000169410 | Tyrosine-protein phosphatase non-receptor type 9 |
| *RABGAP1L* | ENSG00000152061 | Rab GTPase-activating protein 1-like |
| *RALYL* | ENSG00000184672 | RNA-binding Raly-like protein |
| *RAPH1* | ENSG00000173166 | Ras-associated and pleckstrin homology domains-containing protein 1 |
| *RASSF3* | ENSG00000153179 | Ras association domain-containing protein 3 |
| *RBFOX1* | ENSG00000078328 | RNA binding protein fox-1 homolog 1 |
| *RBM20* | ENSG00000203867 | RNA-binding protein 20 |
| *RCAN2* | ENSG00000172348 | Calcipressin-2 |
| *RGS5* | ENSG00000143248 | Regulator of G-protein signaling 5 |
| *RUNX2* | ENSG00000124813 | Runt-related transcription factor 2 |
| *RUSC2* | ENSG00000198853 | AP-4 complex accessory subunit RUSC2 |
| *SGCD* | ENSG00000170624 | Delta-sarcoglycan |
| *SIPA1L2* | ENSG00000116991 | Signal-induced proliferation-associated 1-like protein 2 |
| *SLC20A2* | ENSG00000168575 | Sodium-dependent phosphate transporter 2 |
| *SLC24A2* | ENSG00000155886 | Sodium/potassium/calcium exchanger 2 |
| *SLC35F5* | ENSG00000115084 | Solute carrier family 35 member F5 |
| *SMAD9* | ENSG00000120693 | Mothers against decapentaplegic homolog 9 |
| *SNX25* | ENSG00000109762 | Sorting nexin-25 |
| *SORBS1* | ENSG00000095637 | Sorbin and SH3 domain-containing protein 1 |
| *SORBS2* | ENSG00000154556 | Sorbin and SH3 domain-containing protein 2 |
| *SPAG16* | ENSG00000144451 | Sperm-associated antigen 16 protein |
| *SPOCK1* | ENSG00000152377 | Testican-1 |
| *SUPT3H* | ENSG00000196284 | Transcription initiation protein SPT3 homolog |
| *TAFA1* | ENSG00000183662 | Chemokine-like protein TAFA-1 |
| *TCF7L1* | ENSG00000152284 | Transcription factor 7-like 1 |
| *TCF7L2* | ENSG00000148737 | Transcription factor 7-like 2 |
| *TGFB2* | ENSG00000092969 | Transforming growth factor beta-2 proprotein |
| *THBS4* | ENSG00000113296 | Thrombospondin-4 |
| *THSD7B* | ENSG00000144229 | Thrombospondin type-1 domain-containing protein 7B |
| *TNNI2* | ENSG00000130598 | Troponin I, fast skeletal muscle |
| *TNNC2* | ENSG00000101470 | Troponin C, skeletal muscle |
| *TNNT3* | ENSG00000130595 | Troponin T, fast skeletal muscle |
| *TPM1* | ENSG00000140416 | Tropomyosin alpha-1 chain |
| *TPM3* | ENSG00000143549 | Tropomyosin alpha-3 chain |
| *VWF* | ENSG00000110799 | von Willebrand factor |
| *ZFPM2* | ENSG00000169946 | Zinc finger protein ZFPM2 |

* N/A indicate genes with unknown function
