## Supplementary Table S2 for "Distinct myofibre domains of the human myotendinous junction revealed by single nucleus RNA-seq"

**Supplementary Table S2** - Antibodies for Immunofluorescence

| **Antibodies for staining of tissue cross-sections** | | |
| --- | --- | --- |
| **HGNC gene symbol (protein name)** | **Primary Antibody (Dilution)** | **Secondary Antibody (Dilution)** |
| *ABI3BP* (ABI Family Member 3 Binding Protein, NESHBP, TARSH) | Rabbit IgG; cat. no. NBP2-48811, Novusbio (1:50) | 488 Donkey Anti-Rabbit IgG (Jackson/711-545-152) (1:100) |
| *FRAS1* (Fraser Extracellular Matrix Complex Subunit 1) | Rabbit IgG; cat. no. PA5-62439, Thermofisher (1:10) | 488 Donkey Anti-Rabbit IgG (Jackson/711-545-152) (1:100) |
| *FREM2* (FRAS1 Related Extracellular Matrix 2) | Rabbit IgG; cat. no. NBP1-85049, Novusbio (1:50) | 488 Donkey Anti-Rabbit IgG (Jackson/711-545-152) (1:100) |
| *ABLIM1* (Actin Binding LIM Protein 1) | Rabbit IgG; cat. no. PA5-58607, Thermofisher (1:10) | 488 Donkey Anti-Rabbit IgG (Jackson/711-545-152) (1:100) |
| *ADAMTSL1* (ADAMTS-Like, Punctin) | Rabbit IgG; cat. no. PA5-63442, Thermofisher (1:10) | 488 Donkey Anti-Rabbit IgG (Jackson/711-545-152) (1:100) |
| *BICD1* (BICD Cargo Adaptor 1) | Rabbit IgG; cat. no. NBP1-85843, Novusbio (1:10) | 488 Donkey Anti-Rabbit IgG (Jackson/711-545-152) (1:100) |
| *FHOD3* (Formin Homology 2 Domain Containing 3, Formactin-2) | Rabbit IgG; cat. no. NBP1-83899, Novusbio (1:50) | 488 Donkey Anti-Rabbit IgG (Jackson/711-545-152) (1:100) |
| *CPM* (Carboxypeptidase M) | Rabbit IgG; cat. no. PA5-51730, Thermofisher (1:5) | 488 Donkey Anti-Rabbit IgG (Jackson/711-545-152) (1:50) |
| *COL22A1* (Collagen XXII) | Guinea pig IgG; provided by Manuel Koch (1:500-1:1000) | 680 Donkey Anti-Guinea-pig IgG (Jackson/706-625-148) (1:100-1:250) |
| *NCAM1* (Neural Cell Adhesion Molecule 1) | Mouse IgG1; cat. no. 347740, MY31 BD biosciences (1:100) | 594 Donkey Anti-Mouse IgG (Jackson/715-585-151) (1:250) |
| *COL4A1*/*COL4A2* (Collagen IV) | Mouse IgG1; cat. no. M3F7* DSHB (1:100) | 594 Donkey Anti-Mouse IgG (Jackson/715-585-151) (1:250) |
| *DMD* (Dystrophin) | Mouse IgG1; cat. no. MA513526, Life Technologies (1:500) | 594 Donkey Anti-Mouse IgG (Jackson/715-585-151) (1:250) |
| *DMD* (Dystrophin) | Mouse IgG2b; cat. no. MANDYS8, D8168, Sigma (1:500) | 594 Donkey Anti-Mouse IgG (Jackson/715-585-151) (1:250) |
| *MYH7* (Myosin heavy chain Type I) | Mouse IgG2b; cat. no. BA.D5* DSHB (1:100) | 568 Goat Anti-Mouse IgG2b (Life tec, A-21144) (1:500) |
| Myosin heavy chain (Human fast fibers) | Mouse IgG1; cat. no. A4.74* DSHB (1:50) | 488 Goat Anti-Mouse IgG2 (Life tec, A-21121) (1:500) |
| **Antibodies for staining of single fibres** | | |
| **HGNC gene symbol (protein name)** | **Primary Antibody (Dilution)** | **Secondary Antibody (Dilution)** |
| *ABI3BP* (ABI Family Member 3 Binding Protein, NESHBP, TARSH) | Rabbit IgG; cat. no. NBP2-48811, Novusbio (1:20) | 488 Donkey Anti-Rabbit IgG (Jackson/711-545-152) (1:100) |
| *ABLIM1* (Actin Binding LIM Protein 1) | Rabbit IgG; cat. no. PA5-58607, Thermofisher (1:20) | 488 Donkey Anti-Rabbit IgG (Jackson/711-545-152) (1:100) |
| *DMD* (Dystrophin) | Mouse IgG1; cat. no. MA513526, Life Technologies (1:50) | 594 Donkey Anti-Mouse IgG (Jackson/715-585-151) (1:100) |
| * M3F7 was deposited to the DSHB by Furthmayr, H. (DSHB Hybridoma Product M3F7)  * BA-D5 was deposited to the DSHB by Schiaffino, S. (DSHB Hybridoma Product BA-D5  * A4.74 was deposited to the DSHB by Blau, H.M. (DSHB Hybridoma Product A4.74)   - All samples were fixated for 12 minutes in Histofix after application of the secondary antibody, except ABLIM1 which was fixed for 5 min in 4% PFA prior to incubation with PA. - A 30 minutes blocking step (5% goat serum, 5% donkey serum, 5% BSA in TBS) was included prior to application of the primary antibody when this improved staining quality (ADAMTSL1, BICD1, FHOD3). | | |
